# DNA methylation signatures of early life adversity are exposure-dependent in wild baboons

**DOI:** 10.1101/2023.06.05.542485

**Authors:** Jordan A. Anderson, Dana Lin, Amanda J. Lea, Rachel A. Johnston, Tawni Voyles, Mercy Y. Akinyi, Elizabeth A. Archie, Susan C. Alberts, Jenny Tung

## Abstract

The early life environment can profoundly shape the trajectory of an animal’s life, even years or decades later. One mechanism proposed to contribute to these early life effects is DNA methylation. However, the frequency and functional importance of DNA methylation in shaping early life effects on adult outcomes is poorly understood, especially in natural populations. Here, we integrate prospectively collected data on fitness-associated variation in the early environment with DNA methylation estimates at 477,270 CpG sites in 256 wild baboons. We find highly heterogeneous relationships between the early life environment and DNA methylation in adulthood: aspects of the environment linked to resource limitation (e.g., low-quality habitat, early life drought) are associated with many more CpG sites than other types of environmental stressors (e.g., low maternal social status). Sites associated with early resource limitation are enriched in gene bodies and putative enhancers, suggesting they are functionally relevant. Indeed, by deploying a baboon-specific, massively parallel reporter assay, we show that a subset of windows containing these sites are capable of regulatory activity, and that, for 88% of early drought-associated sites in these regulatory windows, enhancer activity is DNA methylation-dependent. Together, our results support the idea that DNA methylation patterns contain a persistent signature of the early life environment. However, they also indicate that not all environmental exposures leave an equivalent mark and suggest that socioenvironmental variation at the time of sampling is more likely to be functionally important. Thus, multiple mechanisms must converge to explain early life effects on fitness-related traits.

**Significance statement:** The environment animals face when young can affect how they function throughout life. Long-lasting changes in DNA methylation—a chemical mark deposited on DNA that can affect gene activity—have been hypothesized to contribute to early life effects. But evidence for persistent, early environment-associated differences in DNA methylation is lacking in wild animals. Here, we show that early life adversity in wild baboons predicts DNA methylation levels in adulthood, especially for animals born in low resource environments and drought conditions. We also show that some of the changes we observe in DNA methylation have the capacity to influence gene activity levels. Together, our results support the idea that early experiences can become biologically embedded in the genomes of wild animals.

## Introduction

Environmental adversity is a key predictor of morbidity, mortality, and Darwinian fitness in animals. In some cases, these effects are immediate. However, in long-lived species, exposure to adversity can be temporally separated from its outcomes later in life (1), creating lagged associations between environmental experience and trait variation. In humans, for example, adverse childhood experiences predict elevated disease risk and years of lost life many decades later (2, 3). Similarly, in natural baboon, hyena, and bighorn sheep populations, individuals exposed to social, ecological, or physical adversity in early life often survive to adulthood, but on average live shorter adult lives (4–6). Experimental studies in rodents and nonhuman primates show that these lagged effects can reflect causal relationships (7–10). For example, captive rhesus macaques separated from their mothers soon after birth exhibit higher rates of illness and stereotyped behavior later in life, and the effect of maternal separation can spill over to a third generation via its effects on parenting behavior (11, 12).

An animal’s past environments can therefore shape its phenotype long after those environments change, even if conditions improve (13, 14). These observations are likely to be explained, at least in part, by the process of “biological embedding”, which posits that differences in life experience produce stable, systematically different biological states that have the capacity to influence physiology, fertility, or survival across the life course (15). Multiple mechanisms have been proposed to mediate the embedding process, including changes in neural connectivity, HPA axis signaling, and cell type composition (15, 16). At the molecular level, the majority of research has focused on environmentally responsive changes to the epigenome, especially those mediated by DNA methylation: the covalent addition of methyl groups to DNA, which, in vertebrates, occurs primarily at CpG motifs (15, 17–19). Patterns of DNA methylation are largely laid down *in utero* and during the first years of life (i.e., during cellular differentiation and tissue formation) and they can be highly sensitive to environmental conditions during this time (20). However, changes in DNA methylation also occur in response to environmental stimuli later in life, including pathogen exposure, metabolic stress, and glucocorticoid signaling (21–24). Because DNA methylation marks can remain stable across cell divisions (25), they provide a plausible route for encoding a memory of past events in the genome. And because DNA methylation can sometimes—although not always—affect downstream gene expression (26–28), such changes could potentially account for trait consequences at the whole organism level.

For DNA methylation to explain lasting effects of environmental experience, at least two requirements must be met. First, variation in DNA methylation must be linked to the environmental exposure of interest, ideally in a manner that excludes confounding by third variable effects. Second, DNA methylation levels must have the capacity to influence downstream phenotypes, most likely through an initial effect on gene expression. Although often assumed in studies of biological embedding, this relationship is not assured: many CpG sites in mammalian genomes are located outside of known regulatory elements or in inactive heterochromatin (18, 27). Additionally, targeted manipulation of DNA methylation levels using epigenome editing or reporter assays shows that methylation-dependent changes to gene regulation are locus-dependent, and sometimes undetectable (28–30), but see also (31). For example, in massively parallel reporter assays testing the regulatory capacity of many loci simultaneously, only a small fraction of tested regions influenced gene regulation in the human genome (29, 32), and only a minority exhibited significantly altered activity as a function of methylation state (29).Thus, candidate CpG sites involved in biological embedding need to be empirically tested before their capacity to affect downstream traits is assumed (17, 33).

In mammals, including humans, evidence of DNA methylation-mediated embedding in natural populations remains limited. In humans, most work has focused on identifying associations between early life experience and DNA methylation levels in samples collected in adulthood (34–36). For example, DNA methylation levels in the blood of individuals exposed *in utero* to the Dutch hunger winter (a period of extreme caloric restriction induced by a German blockade during World War II: (37)) differ from unexposed individuals near genes involved in growth and metabolism (38). Similarly, people born in rural Gambia during the wet season (a period of relatively high malarial burden and low food availability) exhibit differences in DNA methylation—measured nearly a decade later—compared to those born in the dry season (39). However, large cohort studies that focus on the typical spectrum of variation in developed nations often find relatively few associations between early adversity and DNA methylation, especially after controlling for confounding factors (e.g., smoking behavior) that also vary as a function of early adversity (34–36, 40). Meanwhile, in natural animal populations, studies of biological embedding via DNA methylation remain rare, power-limited, and focused on global rather than site-specific measures of DNA methylation levels (41, 42). For example, higher levels of maternal care and subadult social connectedness predict higher global DNA methylation levels in wild spotted hyenas, but the individual regulatory elements, genes, and pathways that drive this observation are unknown (42, 43). Finally, in both human and nonhuman animal studies, analyses typically stop after identifying putative early life-DNA methylation associations. Without testing the functional consequences of DNA methylation at early environment-associated sites, the importance of DNA methylation in biological embedding remains unclear.

To address this gap, we investigated locus-specific associations between DNA methylation and major sources of early life adversity in a longitudinally studied population of wild baboons living in the Amboseli ecosystem of Kenya (n=256 individuals; 115 male, 141 female) (44). We combined DNA methylation data on nearly half a million CpG sites genome-wide with five decades of ecological, behavioral, and life history data for individually recognized baboons followed across the life course. Importantly, strong early life effects on physiology, fertility, and survival are well-established for this population and for baboons and nonhuman primates more generally (5, 45–49). In Amboseli, female baboons who experience high levels of early life adversity die at substantially younger ages, on average, than those who experience little to no early adversity (5). These females also have elevated glucocorticoids in adulthood (50) and weaker social bonds (5), and their offspring are less likely to survive to adulthood (45).

In addition to five sources of early adversity that have been extensively studied in the Amboseli baboons (5, 45, 51), we also investigated the effect of habitat quality, a primary driver of resource availability in our population. In particular, large differences in habitat quality differentiate study subjects who were born early in the long-term study period (when the two original study groups shifted their home ranges to a new part of the study site) from those born after the home range shift. This shift was precipitated by a rapid die-off of fever trees (*Vachelia xanthophloea*), a major source of food and protection from predators, in the pre-shift habitat.

After the home range shift, female baboons experienced shorter inter-birth intervals, began reaching reproductive maturation earlier, and exhibited improved infant survival rates (52, 53), in support of an improved resource base. We therefore included habitat quality at birth (pre-shift or post-shift: Fig. S1) as another source of early life disadvantage.

By integrating our measures of early life adversity with genomic data on DNA methylation and gene expression, we were able to pursue four major goals. First, we tested for a signature of early life adversity on DNA methylation levels in blood, including how sources of early adversity that differentiate animals within the same group interact with overall habitat quality in early life. To place our results in context, we compared the signature of early adversity to those of dominance rank (i.e., social status) at the time of sampling, an important predictor of gene regulation in the Amboseli baboons and other mammals (54–57), and of age, a major predictor of DNA methylation across mammals (54, 58, 59). Second, we investigated how the DNA methylation signatures of distinct environmental variables are distributed across the genome and whether they overlap with one another. Importantly, major sources of early life adversity in the Amboseli baboons are not well-correlated with each other, and early life experience is also usually uncorrelated, or weakly correlated, with the adult environment (Fig. S2) (5, 45). These features of our study system enabled us to disentangle the DNA methylation signatures associated with distinct environmental exposures, a perennial challenge in humans (3). Third, we asked whether the signature of habitat quality in early life weakens with temporal distance from early life, as predicted if experiences in adulthood also modify the epigenome. Finally, we coupled experimental *in vitro* evidence from a massively parallel reporter assay, mSTARR-seq (29) and *in vivo* evidence from gene expression samples from the same population (60) to investigate whether, when, and how often DNA methylation levels at environment-associated CpG sites are likely to be functionally relevant for gene regulation in blood.

## Results

### DNA methylation levels are associated with environmental variation in early life and adulthood

To investigate the signature of environmental variation on the baboon DNA methylome, we used reduced-representation bisulfite sequencing (RRBS (61, 62)) to profile DNA methylation in blood for 477,270 CpG sites in the baboon genome, in 256 unique individuals (115 males, 141 females). For 37 individuals, we profiled repeated, longitudinally collected samples (2-3 samples per individual), for a total of n=295 samples (Table S1).

For each CpG site separately, we first modeled DNA methylation levels as a function of habitat quality at birth, cumulative early life adversity, and age and ordinal dominance rank at time of sampling, using the binomial mixed effects model implemented in *MACAU* (Model 1; Fig. 1A; see SI methods for model details) (58). We quantified habitat quality at birth as a simple binary variable indicating whether each study subject was born before or after the home range shift described above (N=57 low quality individuals). We treated habitat quality at birth separately from cumulative early adversity because of its nature as a strong cohort effect characterized by two distinct time periods, rather than a set of conditions that vary across individuals living at the same time and place. We considered five sources of early adversity as components of the cumulative early adversity measure: drought, maternal loss, large group size, the presence of a close-in-age younger sibling, and low maternal dominance rank, which collectively predict both reduced survival and reduced offspring survival (5, 45) (see also Methods). We estimated dominance rank effects for each sex separately (by nesting rank within sex), as male and female ranks depend on different traits for each sex (i.e., kinship in females and physical condition in males). Further, the hierarchies for each sex are separately estimated, have sex-specific implications, and have sex-specific associations with gene expression (44, 60, 63–66).

**Fig. 1.**
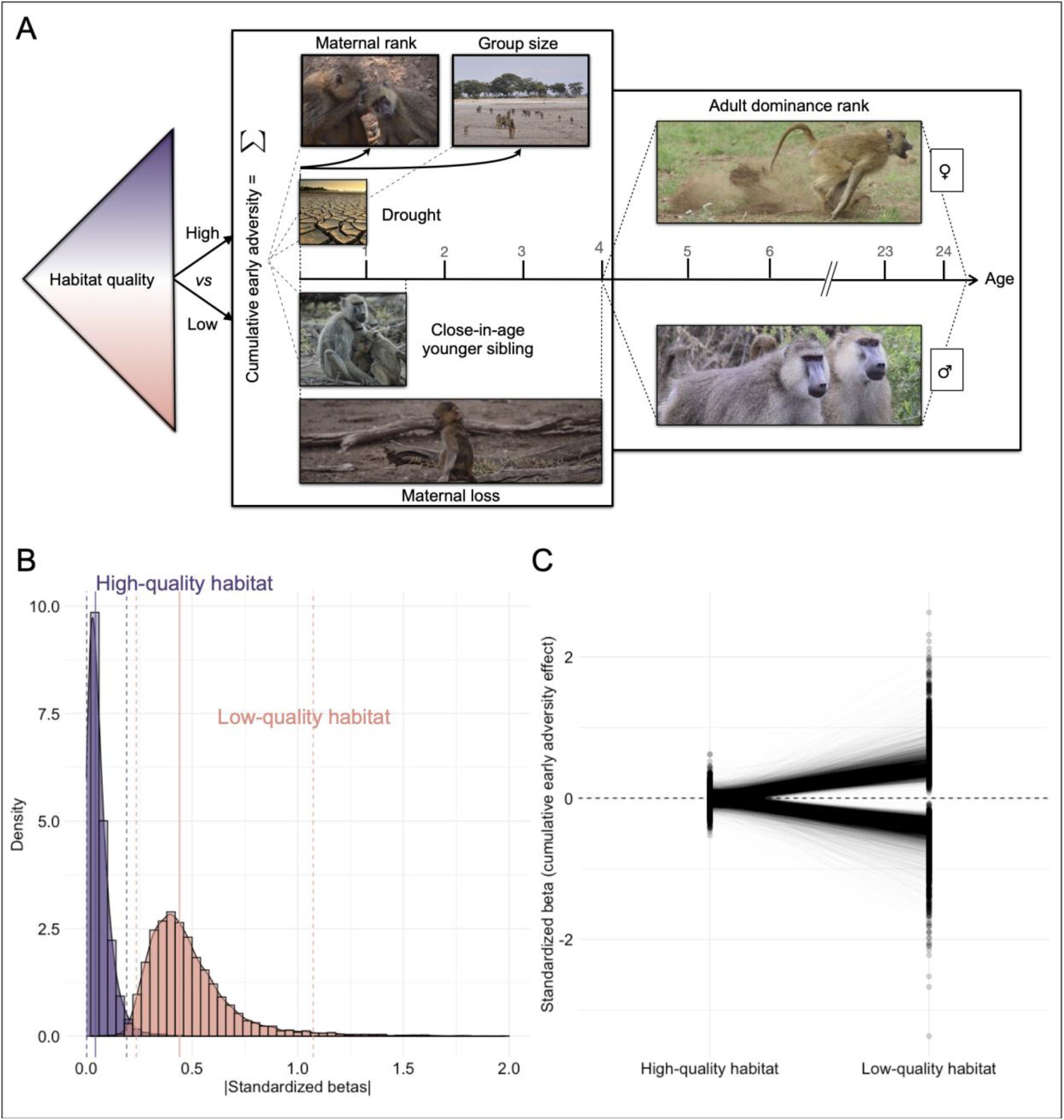
Socioenvironmental predictors of DNA methylation depend on early life habitat quality. (A) Study design: we investigated (i) habitat quality at birth (lefthand triangle: high/post-shift versus low/pre-shift), (ii) cumulative exposure to each of five individual sources of early adversity (lefthand box), and the effects of (iii) age and (iv) dominance rank at the time of sample collection (right hand box). (B) The absolute value of cumulative early adversity effects estimated for individuals born in high-quality habitat (purple) versus those born in low-quality habitat (peach) for sites passing a 20% FDR in one or both conditions (n=12,872 CpG sites; Model 2). Solid and dashed lines show the mean and 95% intervals, respectively, for each distribution. (C) Standardized betas, from Model 2, comparing the effect of cumulative early adversity for individuals born in low-versus high-quality habitats, across the same set of sites (n=12,872). Each line connects the two effect sizes for one CpG site (one effect size estimate from samples of individuals born in the high-quality habitats and the second estimated for those born in low-quality habitats).

In Model 1, the strongest predictors of DNA methylation in adulthood were habitat quality at birth, male dominance rank at sample collection, and age at sample collection. The relationship between habitat quality at birth and DNA methylation was striking (3,296 habitat quality-associated sites, FDR=10%; Table S2A). Consistent with effects of dominance rank on other aspects of gene regulation (60, 66), associations between male dominance rank and DNA methylation were also widespread (n=3,736 sites, 10% FDR), in contrast to a weaker relationship with female dominance rank (n=4 sites). Age strongly predicted DNA methylation across the genome (n=169,439 age-associated sites), with a bias, as reported in other studies (67, 68), to increases in DNA methylation with age in CpG islands (65%) and decreases in DNA methylation with age in most other regions of the genome (79%). In contrast to these three effects, we observed no significant associations (10% FDR) between DNA methylation and cumulative early adversity.

Our results for Model 1 suggest that habitat quality in early life is particularly important in the lives of baboons and could moderate the effect of other sources of early adversity on DNA methylation. To test this possibility, we re-ran our analyses, but in this case tested for the effects of cumulative early adversity experienced in the high-quality habitat and low-quality habitat separately (i.e., by nesting cumulative early adversity within habitat quality; Model 2). To maximize power, we also included individuals for whom early adversity data were available, but dominance rank data were missing because of observational gaps for males. This model not only strengthens the evidence for a main effect of habitat quality (25,509 habitat quality-associated sites; 10% FDR), but reveals an interaction with cumulative adversity: 2,856 sites are associated with cumulative adversity for baboons born in low-quality habitat (10% FDR), while none are significantly associated with cumulative adversity in baboons born in high-quality habitat (Fig. 1B, 1C; Table S2B). Notably, only 64 of 295 samples derive from low-quality habitat individuals, suggesting that the greater magnitude of effects in low-quality habitat are not driven by greater power. Among the significant sites identified in samples from individuals born in low-quality habitat, the effect sizes for cumulative adversity in the low-quality habitat are uncorrelated with the effect sizes for cumulative adversity in high-quality habitat (p=0.838) but positively correlated with the effect sizes for habitat quality itself (R=0.508, p<1 × 10^−10^), suggesting that the effect of cumulative adversity is amplified by exposure to ecologically challenging conditions (and vice-versa). Importantly, cumulative adversity scores do not differ between animals born in low-quality and high-quality habitats (Wilcoxon rank-sum test p=0.843).

To investigate whether different components of the cumulative adversity score contribute differently to these effects, we then ran a third model (Model 3) to evaluate each of the five individual sources of early adversity, nested within habitat quality (all other biological and technical covariates remained the same as in Model 2). Among the individual sources of adversity we considered, early life drought most clearly predicted variation in DNA methylation across the genome, especially for individuals born in low-quality habitat (25,355 sites; Fig. 2A,2B). We also identified detectable, but less common signatures of maternal loss (4,893 sites), large group size (3,124 sites), low maternal rank (730 sites), and the presence of a close-in-age younger sibling (619 sites). In contrast, none of the individual sources of early adversity were robust predictors of DNA methylation for individuals born in the high-quality habitat (≤5 sites associated with any individual predictor at 10% FDR; Fig. 2C; Table S2C).

**Fig. 2.**
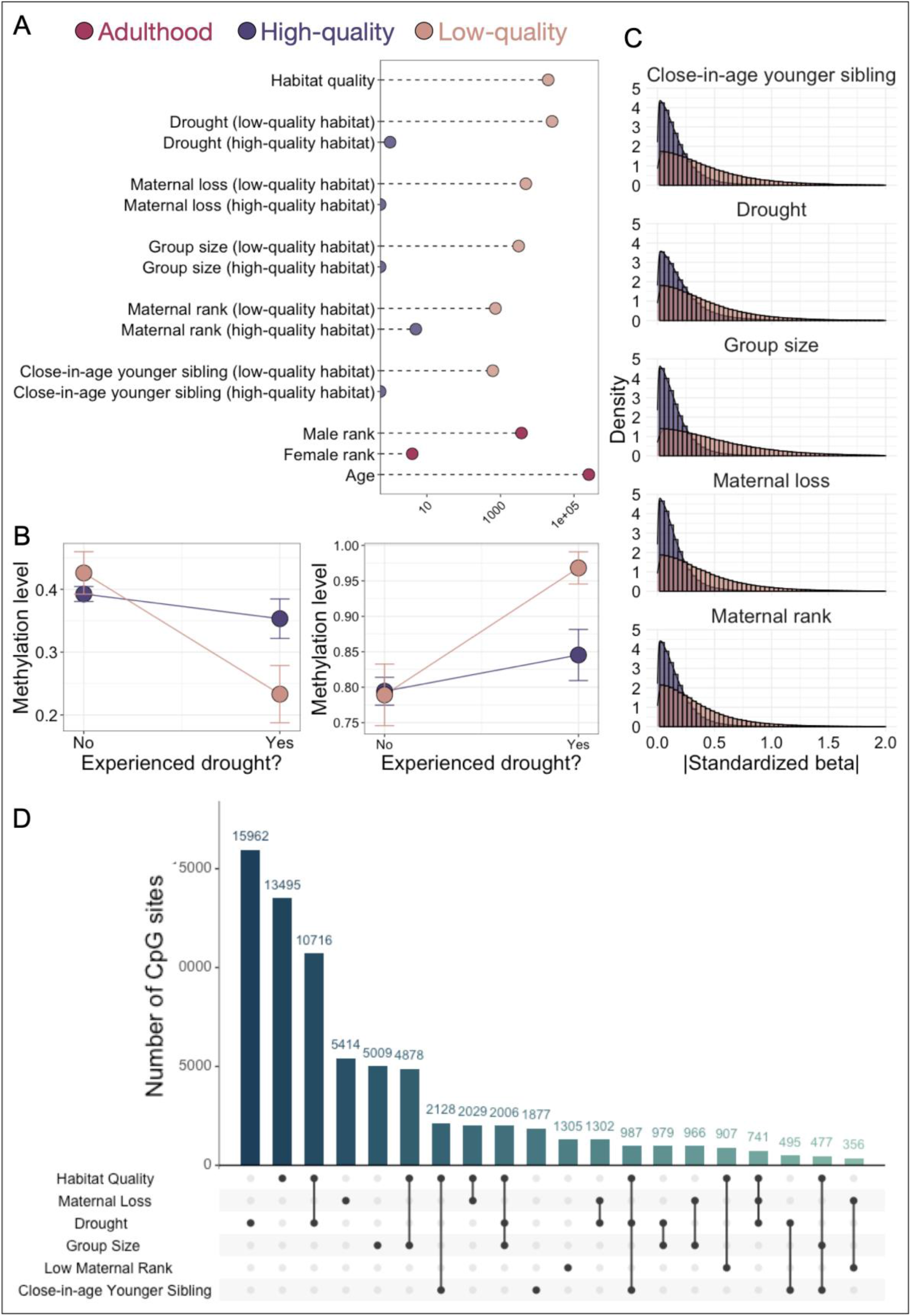
Early life adversity is associated with DNA methylation in adulthood for baboons born in low-quality habitat. (A) The number of CpG sites associated with each tested predictor (<10% FDR) in Model 3. The x-axis is shown on a log10 scale. (B) Reaction norms for two example CpG sites (chr12_111013997 and chr11_430191) that were significantly associated with early life drought, but only for baboons born in low-quality habitat (peach; 10% FDR). Colored bars indicate standard errors. (C) Distributions of the absolute value of standardized effect sizes across tested sites for each of five individual-level sources of early adversity. In all cases, effect sizes are systematically larger for individuals born into low-quality habitat (peach) environments than those born into high-quality environments (purple). (D) UpSet plot of the number of CpG sites associated with habitat quality, each individual source of adversity (within low-quality habitat), and their overlap. Each bar represents the number of sites associated with the source(s) of adversity indicated in the matrix beneath the bar graph. To avoid calling sites “unique” due to small differences in FDR values, overlaps show sites that are significant at a 10% FDR threshold for at least one predictor variable and *≤*20% FDR for the other predictor variable(s).

### The genomic distribution of environmental predictors of DNA methylation

Our models indicate that some early life experiences are linked to more pronounced DNA methylation signatures than others. Drought in particular, which is one of the least predictable environmental exposures in Amboseli, is associated with an order of magnitude more CpG sites than maternal rank or group size, the next most common effects. Notably, early life and rainfall at the time of sampling are only weakly correlated in our dataset, supporting the idea that our observations capture long-term early life effects (Fig. S3). To investigate whether these signatures are unique to specific early life experiences or reflect a general signature of stress and adversity (perhaps scaled to the magnitude of the stressor), we therefore tested for overlap between the sets of sites linked to each of the five individual-level predictors and to habitat quality based on results from Model 3.

Our results support a generalized rather than an exposure-specific signature (Fig. 2D).

Specifically, among sources of early adversity with a substantial number of associated CpG sites (habitat quality, drought, maternal loss, and group size), sites associated with one early life exposure are 1.04 – 8.6-fold more likely to be associated with a second early life exposure (p < 1 × 10^−10^ for 4/6 comparisons). Habitat quality and drought (in samples from individuals born in low-quality habitat) show a particularly striking pattern of overlap: 4,038 CpG sites are significantly associated with both predictors (log_2_(OR)=2.23, p< 1 × 10^−10^), and 4,030 of these cases (99.8%) are directionally concordant, such that exposure to low habitat quality in early life and exposure to drought predict the same direction of effect.

Comparing these findings to the signature of male dominance rank shows that overlap in sensitivity to the environment is not specific to early life effects (note that we focused on male rank here because significant associations with female rank are far less common). Male rank-associated sites are 11.21 times more likely to be associated with drought than background expectations and 2.43 times more likely to be associated with habitat quality (both p < 1 × 10^−10^). In these cases, dominance rank effects tend to have directionally opposite effects to habitat quality and drought (log_2_(OR)=-4.06 for overlap with habitat quality; the odds ratio could not be estimated for the overlap with drought because there was no overlap in the direction of effects). Consequently, sites that are more highly methylated in high-ranking males also tend to be more highly methylated for baboons of both sexes who were born in poor quality habitat and exposed to drought within that habitat.

In contrast to male rank-associated patterns of DNA methylation, age effects on DNA methylation only modestly overlap with drought effects and habitat quality (log_2_(OR)=0.16 and 0.43, both p<10^−10^) and do not overlap with male rank effects at all (log_2_(OR)=0.045, p=0.35) (Fig S4). These results suggest that despite a shared epigenetic signature of at least some types of early and adult experience (with variation in the magnitude of the effect), the effects of age are distinct. To test this hypothesis further, we investigated how CpG sites related to age versus socioenvironmental variables are distributed across promoters, gene bodies, CpG islands and shores, putative enhancer elements, and unannotated regions. We focused on the four variables with the strongest DNA methylation signatures: age, habitat quality in early life, drought (in the low-quality habitat), and male dominance rank. Our results highlight two patterns (Fig. 3A). First, drought and male dominance rank-associated sites are systematically enriched in functionally important regions of the genome, especially gene bodies (log_2_(OR)=0.25 and 0.72, respectively) and putative enhancer elements (log_2_(OR)=0.52 and 0.99), but depleted in unannotated regions (log_2_(OR)=-0.13 and -0.36) of the genome (all p<1×10^−7^; Table S3).

**Fig. 3.**
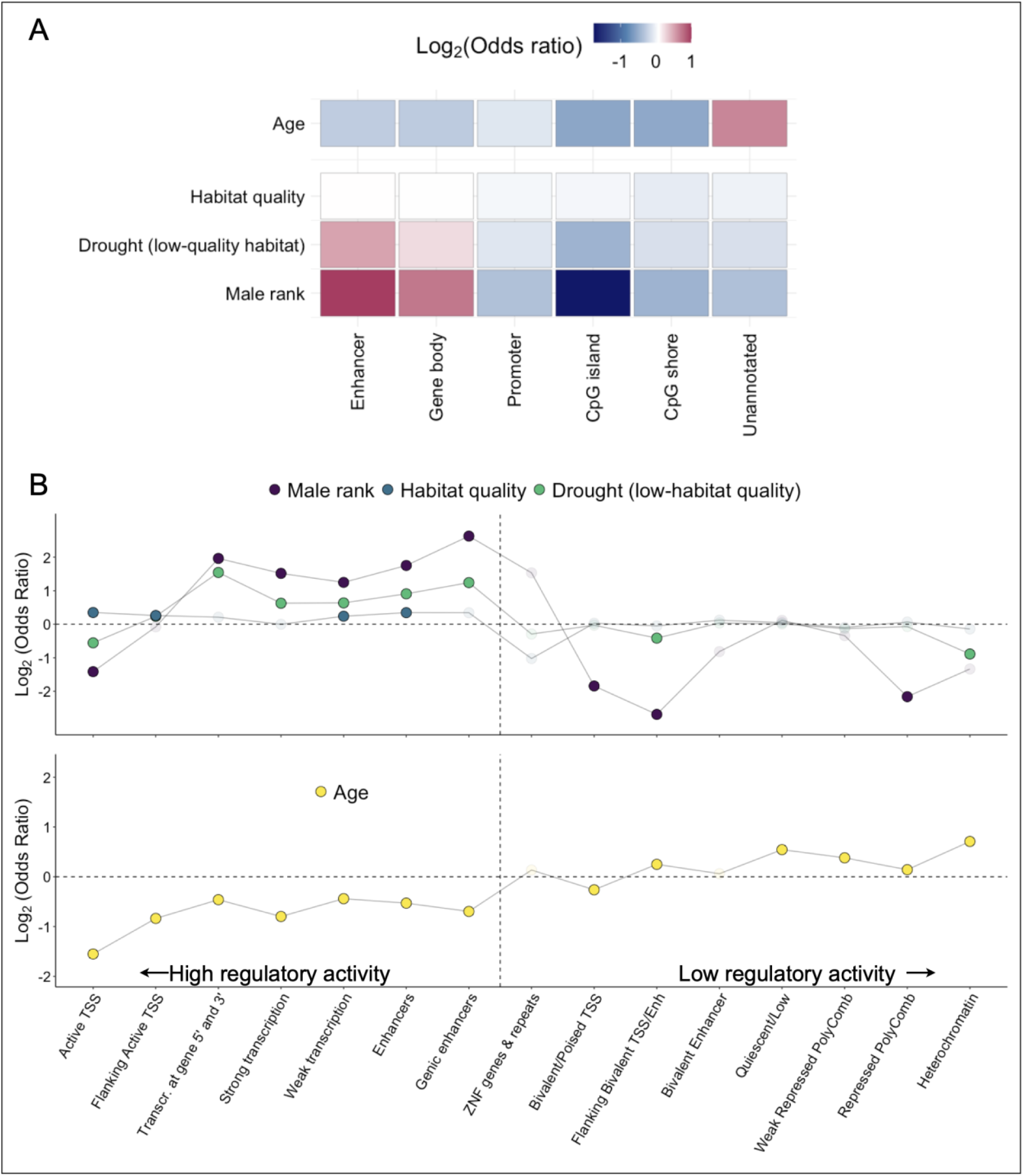
Genomic distribution of CpG sites associated with age, rank, and early life adversity. (A) Enrichment of the top four predictors of DNA methylation levels in functional compartments across the genome. Color indicates log2(Odds Ratio) from a Fisher’s exact test, with the brightest colors indicating highest and lowest odds ratios. (B) Enrichment of the same four sets of age, rank, or early environment-associated CpG sites, across 15 distinct chromatin states, based on annotation in human peripheral blood mononuclear cells with coordinates lifted over to *Panubis1*.*0*. States are ordered roughly by their association with active gene regulation, from left (active) to right (repressed/quiescent). Opaque dots indicate p<0.05 for enrichment based on Fisher’s exact test.

Second, and in contrast, age-associated sites are 1.57-fold more likely to occur in unannotated regions of the genome than expected by chance, but are depleted in enhancers (log_2_(OR)=-0.27) and gene bodies (log_2_(OR)=-0.28, all p<1×10^−10^). Notably, habitat quality-associated sites, which are much more widely distributed in the genome than rank- or drought-associated sites, follow an intermediate pattern: they are less common in unannotated regions than age-associated sites but are not strongly enriched for gene bodies or enhancers.

A similar two-class pattern is observable based on enrichment within chromatin state annotations (i.e., predictions of the function of different regions of the genome based on the presence of 5 epigenetic marks: H3K4me3, H3K4me1, H3K36me3, H3K27me3, H3K9me3). Here, we lifted over chromatin state coordinates for human peripheral blood mononuclear cells to the baboon genome, *Panubis1*.*0* (27, 69). Early life drought and dominance rank are again enriched in regions of the genome marked for regulatory activity, such as enhancer elements (log_2_(OR)=0.91 and 1.75 respectively, both p<10^−10^), and transcriptional activity (log_2_(OR)=0.62 and 1.52 respectively, both p<10^−10^), but depleted in repressed and silenced regions such as heterochromatin (log_2_(OR)=-0.89 and -1.32, p= 4.4 × 10^−5^ and 0.055) and weakly repressed, polycomb-marked DNA (log_2_(OR)=-0.13 and -0.34, p=0.03 and 0.016; Fig. 3B top; Table S3). Age-associated sites show the opposite pattern (Fig. 3B bottom).

### The DNA methylation signature of early life habitat quality attenuates over time

Although the individuals in our data set were predominantly adults, individuals exposed to poor habitat quality were sampled at a range of ages (range=2.5-26.3 years). We took advantage of this variation to test whether the signature of early life adversity attenuates over time, resulting in weaker signatures with longer times from exposure. To do so, we focused on habitat quality, the strongest early life effect we observed in our data. We first built an elastic net model to ask whether early life exposure to low-quality habitat (a binary variable indicating whether the subject was born before or after the home range shifts) is predictable based on DNA methylation levels sampled in adulthood (70).

We found that an elastic net model achieves high accuracy in our sample (AUC=0.92 based on leave-one-out cross-validation; Fig. 4A-B). However, among animals born in low-quality habitats, the ability of the model to correctly and confidently predict habitat quality in early life depends on the time elapsed between the habitat shift and blood sample collection (linear model p=0.0084; Fig. 4C), but not on cumulative amount of time spent in low quality habitat (linear model p=0.279). Animal age does not predict overall habitat quality, so we are not indirectly capture habitat quality through age of individuals (linear model p=0.80). Specifically, animals who had spent more time in high-quality habitat prior to sampling were less confidently predicted to be born in low-quality habitat than those who experienced it more recently. This result suggests that, although DNA methylation signatures of early adversity can persist for years in baboons, they also decay over time or are overwritten by later life experience.

**Fig. 4.**
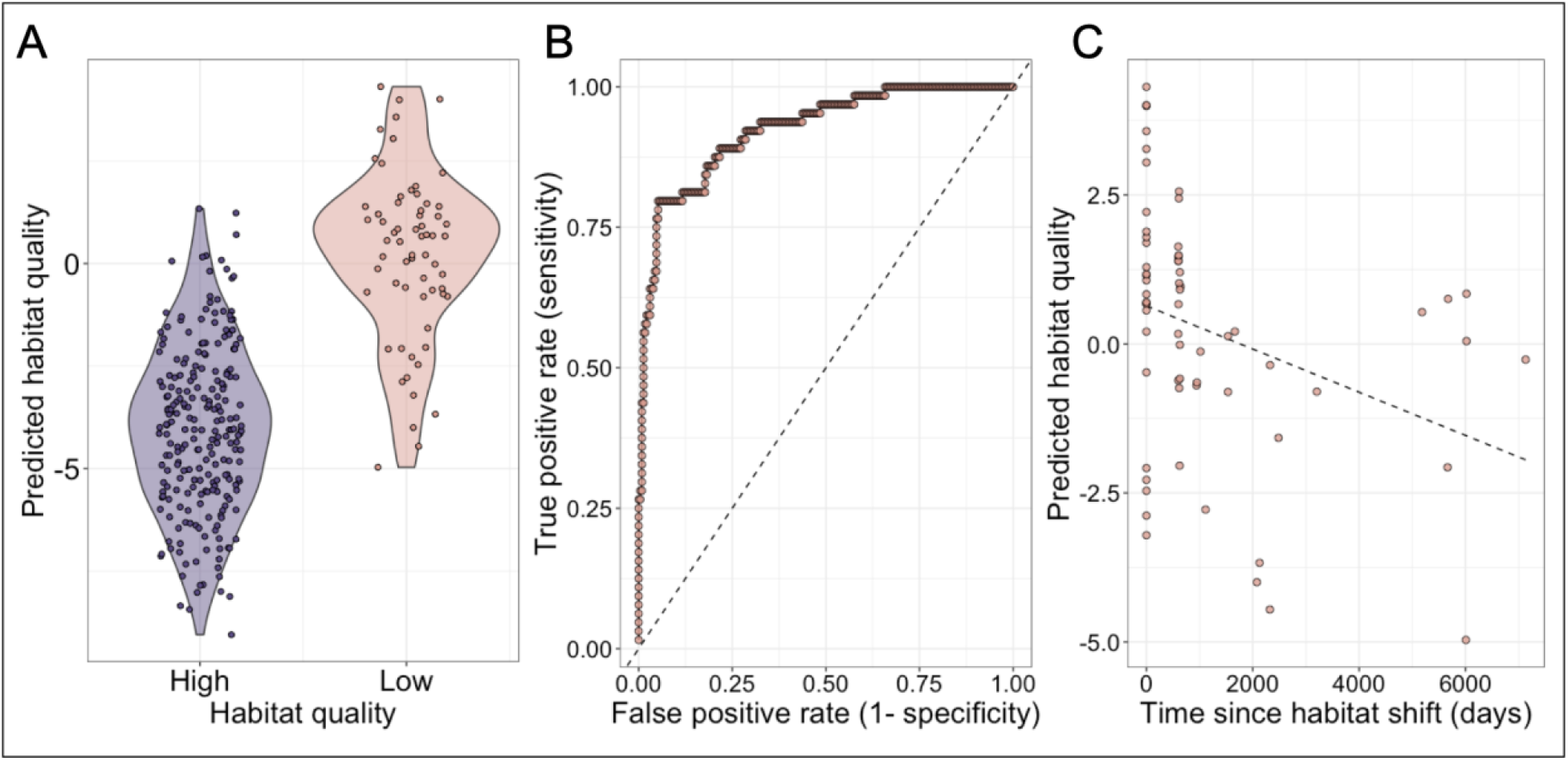
Early life habitat quality can be accurately predicted from DNA methylation, but this signal attenuates over time. (A) Known early life habitat quality (x-axis) versus predicted early life habitat quality from an elastic net regularization model (y-axis). More negative values correspond to cases in which the model predicted that the individual was born in high-quality habitat (the post-habitat shift environment); more positive values correspond to cases in which the model predicted that the individual was born in low-quality habitat (the pre-shift environment). (B) Receiver operating characteristic (ROC) curve for early life habitat quality predictions (AUC=0.926; dashed line denotes the y=x line). (C) Predicted habitat quality (y-axis) versus the time since habitat shift in days (x-axis) for animals born in low quality habitat (linear model p=0.0084). 0 days since habitat shift indicates a sample from an animal still in the low-quality environment.

### Evidence for the functional importance of environment-associated DNA methylation variation

The distribution of environment-associated CpG sites in loci related to transcription and active gene regulation suggests that some subset of these sites have the capacity to causally influence gene expression. To formally test this hypothesis, we performed a massively parallel reporter assay (MPRA), mSTARR-seq, designed to both identify loci capable of regulatory activity *in vitro* and quantify the effects of differential methylation on the magnitude of this activity (Fig 4A; Table S4) (29). mSTARR-seq tests a sequence fragment’s ability to drive gene expression in a self-transcribing plasmid, in hundreds of thousands of genomic fragments simultaneously. Fragments capable of driving their own transcription have enhancer-like activity *in vitro*. Since the plasmid backbone is devoid of CpG sites, inserted fragments containing CpG sites in their sequence can be tested in either a fully CpG methylated or fully unmethylated state to investigate whether enhancer activity can be modified by changes in DNA methylation alone.

We performed mSTARR-seq using a mechanically fragmented and restriction enzyme-digested library of baboon DNA fragments, transfected into the human K562 cell line (Table S4). K562s are a myelogenous leukemia line that shares properties with several types of peripheral blood mononuclear cells and are therefore often used in studies of immune variation.

Importantly, mSTARR-seq has also been extensively optimized in K562 cells (29). Following quality control, we were able to test for regulatory activity in 252,463 500-base pair windows across the baboon genome (4.4% of the genome), of which 32,634 contained tested CpG sites in the Amboseli baboon data set. Among these 32,634 windows, we identified 492 windows (1.5% of those tested, using a 10% FDR threshold; Table S5) capable of enhancer activity in either an unmethylated state, a fully methylated state, or both (similar to estimates from (29)).

As expected, experimentally identified regulatory regions were strongly enriched in predicted strong enhancers (based on chromHMM annotations: log_2_(OR)=2.50. p<10^−10^) and depleted in insulators (log_2_(OR)=-2.32 p=2 × 10^−9^) and repressed regions (log_2_(OR)=-0.94, p=5.8 × 10^−18^; Fig 4B; Table S6). Among the 492 regulatory windows overlapping our tested sites, 86% also exhibited methylation-dependent activity, where the capacity to drive transcription differs depending upon whether CpG sites are methylated or not (Fig 5A). 94% of methylation-dependent regions exhibited reduced activity in the methylated state compared to the unmethylated state, consistent with a general role for DNA methylation in repressing gene regulation.

**Fig. 5.**
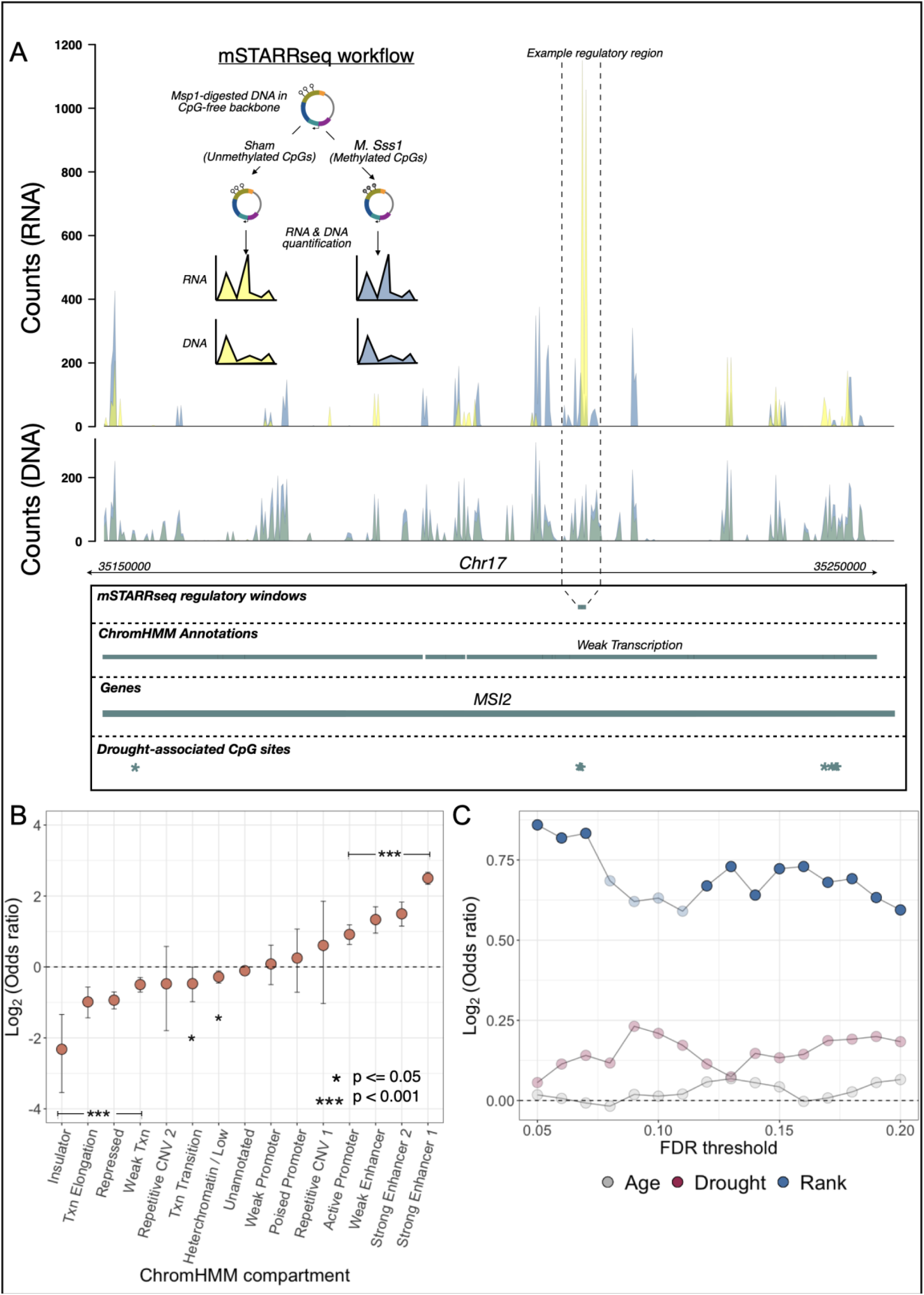
CpG sites associated with drought and male dominance rank are enriched in functional regions of the genome based on a high-throughput reporter assay. (A) Workflow for the mSTARR-seq experiment and an example of read pileups at a regulatory window that exhibits methylation-dependent regulatory activity and overlaps a drought-associated CpG site in the observational data from Amboseli. Summed read counts are shown for methylated (blue) and unmethylated (yellow) experimental replicates. In the highlighted methylation-dependent regulatory region, unmethylated treatments drive substantial expression (yellow RNA counts) compared to methylated treatments (blue RNA counts), even though the amount of input DNA (overlapping yellow and blue DNA counts) was near-identical across treatments. (B) Enrichment of regulatory regions from mSTARR-seq across 15 chromatin states lifted over to the baboon genome from human peripheral blood mononuclear cells (27). Regions with empirically identified regulatory activity are enriched in regions orthologous to putative enhancer and promoter regions in human PBMCs, and depleted in states associated with regulatory quiescence/repression. (C) Enrichment statistics for male dominance rank-(blue), drought-(red), and age-associated CpGs (gray) in regions capable of regulatory activity in mSTARR-seq. The x-axis shows the FDR threshold for identifying age, drought, or rank-associated CpG sites; the y-axis shows the log2(OR) for enrichment in mSTARR-seq putative regulatory elements (all identified at FDR = 10%). Opaque points indicate significant FET enrichment (p<0.05).

For most discovery thresholds between 5% and 20% FDR, male dominance rank-associated CpG sites are found in mSTARR-seq-identified regulatory windows more often than chance, such that most discovery threshold-predictor variable combinations reach statistical significance (Fig. 5C). For example, 40 dominance rank-associated sites (FDR = 20%; 2.2% of significant sites tested) fall in regions of the genome capable of behaving like enhancer elements (log_2_(OR)=0.59 FET p=0.014; note that promoter regions often also exhibit enhancer-like activity in massively parallel reporter assays (29)). These sites are also enriched in windows where modifying the DNA methylation level of the tested sequence alters its capacity to drive gene expression (log_2_(OR)=0.71 FET p=0.006). The pattern for drought-associated sites is less clear: while they are not more likely to occur in mSTARR-seq regulatory elements than expected by chance (drought-associated sites at FDR = 20%, n = 196 sites, (log_2_(OR)=0.18 FET p=0.1), they exhibit a modest enrichment for DNA methylation-dependent activity (log_2_(OR)=0.23, FET p=0.06). Age-associated sites again provide a clear contrast, with no evidence of enrichment of such sites among mSTARR-seq-identified regulatory windows (log_2_(OR)=0.065, p=0.45).

These results suggest that the associations detected in our field-based sample partially reflect targeted, functionally important changes in the response to the environment, some of which are detectable years to decades post-exposure. If so, environmental effects on DNA methylation should also colocalize with environmental effects on gene expression in the Amboseli baboons. To test this possibility, we drew on RNA-seq gene expression data from white blood cells collected from 2013 – 2018, in which several thousand associations between male rank and gene expression have previously been identified (60, 66) (note that individuals born in low-quality habitat are not well-represented in this data set because the start of collection for gene expression analysis long post-dated the habitat shift). Male rank-associated CpG sites fall closer to, and more often within, genes associated with male dominance rank than they do for the background set of tested genes (Kolmogorov-Smirnov test p=1.81 × 10^−5^). Blood-expressed genes that contain a male rank-associated CpG site are also 1.22-fold more likely to exhibit male rank-associated gene expression levels (p=6.60 × 10^−4^), even though the individuals represented in the gene expression data set and the DNA methylation data set are largely distinct (34 of 115 males in the DNA methylation data set were included in the gene expression data set; 34 of 52 males in the gene expression data set were included in the DNA methylation data set). Finally, rank effects on gene expression are negatively correlated with rank effects on DNA methylation for CpG sites in the same gene (Fig. S5). Thus, if DNA methylation levels are higher in high-ranking males, gene expression levels tend to be lower in high-ranking males, and vice-versa (FET for sign: log_2_(OR)=-1.25 p=8.6 × 10^−9^). As a result, multiple pathways enriched among rank-associated genes based on gene expression are also enriched among genes linked to rank-associated DNA methylation patterns, including interferon alpha signaling, NFkB signaling, and the inflammatory response (all p<0.05; Table S7).

## Discussion

Although early life effects on fitness are documented in many long-lived species, how these effects bridge across time to link the early environment with trait outcomes later in life is not well understood. Here, in support of the biological embedding hypothesis, we find that DNA methylation may serve as a persistent link between some forms of early life adversity and later life phenotypes in wild baboons. We also document a shared fingerprint of early life adversity and male dominance rank (i.e., social status) in adulthood, which is in turn distinct from the much more widespread effects of age. Finally, we leverage *in vitro* experiments and gene expression data from the same population to show that a subset of environment-associated changes in DNA methylation are functionally relevant to gene regulation.

Our results also highlight that not all sources of early adversity—even ones that have substantial effects on fertility and survival—are strong predictors of variation in DNA methylation. For example, maternal loss, the strongest independent predictor of lifespan in female baboons (5), has no detectable relationship with DNA methylation patterns for individuals born into a high-quality habitat, and only a moderate association with DNA methylation for those born into a low-quality habitat. In contrast, the effects of low-quality habitat, and drought exposure within a poor habitat, are widespread. These results support the idea that types of early life adversity that involve resource deprivation may have stronger links to later life DNA methylation patterns than those involving threat (71). Indeed, drought in Amboseli, when yearly rainfall is similar to desert biomes in the American southwest, represents a serious source of resource deprivation (46, 72). Drought in the impoverished habitat pre-range shift, when infant survival rates were 19% lower than in the post-shift high-quality habitat (53), was likely even more challenging. The fact that we were only able to detect drought-associated sites in animals born in the low habitat quality environment therefore suggests that biological embedding via DNA methylation is most pronounced and/or most consistent under conditions of considerable material deprivation. This result may also account for observations in humans, in which DNA methylation associations with early life famine have been discovered more often than associations with early life stressors such as parental loss and poor maternal bonding (35) (38, 39, 73).

A clear implication of our results is that different sources of early life adversity can have compounding effects on DNA methylation. Specifically, all individual-level early life effects we considered were magnified for individuals born into poor early life habitat. This observation suggests that, as reported in studies of adverse childhood experiences, health, and longevity in humans, the effects of combined early adversity can interact to exceed that expected from additive effects (74). We speculate that such interactions are particularly likely to occur for components of the environment that have similar mechanisms of action. Both drought and low-quality habitat, for instance, are costly because they constrain the baboons’ resource base.

Hence, they are likely to affect DNA methylation patterns at a shared set of loci and in a common direction. The baboons’ behavioral choice to shift home ranges may therefore have had long-term ramifications for population variation in both DNA methylation and gene expression profiles.

Our findings also emphasize the importance of explicitly testing for the functional effects of environment-associated DNA methylation and gene regulation. The conventional model for CpG methylation and gene expression, which proposes that DNA methylation causally alters the expression of nearby genes by altering chromatin accessibility and/or transcription factor binding, does not apply to all CpG sites. Indeed, genomic analyses of the response to stimuli show that changes in DNA methylation often occur downstream of transcription factor binding or changes in gene expression (21), rather than the reverse; indeed, changes in DNA methylation have recently been suggested to be dispensable for the function of many enhancer elements (31). For DNA methylation to mediate biological embedding, however, it must play a functional role. And while our results combine with those of others (75) to show that changes in DNA methylation can indeed precede changes in gene regulation—196 drought associated CpG sites identified here fall in regulatory regions with methylation-dependent activity *in vitro*—this pattern is far from universal. For example, in this analysis, roughly 25,000 drought-associated sites either do not fall in regions with enhancer activity in our assay, or are in methylation-insensitive regulatory regions. This observation suggests that many early adversity-associated sites may be functionally silent, exert effects on gene regulation but not via enhancer activity, or have tissue- or environment-specific effects invisible in our single-cell type assay. In either case, empirically testing for the functional consequences of differential methylation can help prioritize environment-associated CpG sites for future work. Such tests should become a standard component of studies of biological embedding.

An important next step will be to identify the factors that mediate and moderate the effects of environmental adversity on DNA methylation, including whether the signatures of adult experiences are more malleable than those that occur during development. For example, previous work has shown that high social status may buffer baboon females from the long-term effects of early life drought on fertility (46), and that strong social bonds and high social status in adulthood can buffer some negative effects of early adversity on survival (76). Whether social status or other advantages in life (e.g., strong social bonds) buffer the relationship between early adversity and DNA methylation remains to be tested. Additionally, DNA methylation levels at many CpG sites have a heritable component (mean h^2^=0.2 in humans: (77)), which our analyses also identifies in the Amboseli baboons (this study: mean h^2^=0.28 ? 0.2 s.d.). Whether genetic variants associated with DNA methylation levels (i.e., methylation quantitative trait loci, or meQTL) co-occur or interact with the effects of early adversity is a natural question to address in future work. Finally, although our results suggest that a subset of early adversity-associated sites have the capacity to also influence gene regulation, whether and how these effects influence organism-level physiology, health, and survival remains a puzzle. Investigating the role of differential methylation at such sites for shaping the molecular response to pathogens, nutrient availability, or hormonal signals of stress (as in (78)), may help resolve this open question.

## Materials and Methods

Study subjects were 256 adult baboons (115 males and 141 females) living in one of the 25 study groups observed by the Amboseli Baboon Research Project (ABRP) between 1979-2018 (Table S1). In all cases, blood samples were obtained via brief anesthetization of each study subject during periodic darting efforts, in which a Telazol-loaded dart was delivered via a handheld blowgun (60, 66, 79, 80). Methylation levels were measured using single or double digest reduced representation bisulfite sequencing of DNA extracted from whole blood. Reads were mapped to the *Panubis1*.*0* genome (GCA_008728515.1), and CpG sites with low coverage or that were constitutively hypo/hyper-methylated were removed, leaving 477,270 sites for downstream analyses.

Measures of early life adversity were prospectively and directly observed through longitudinal monitoring of the population. Similar to (5), and following (45), we quantified cumulative early adversity as the sum of exposures to five major sources of environmental adversity in early life: low maternal dominance rank (lowest quartile of ordinal ranks in the population, where higher numbers correspond to lower social status), social group size at birth (highest quartile) as an index of resource competition, drought in the first year of life (<200 mm of total rainfall), the presence of a close-in-age younger sibling (live birth within 1.5 years of the focal individual, approximately the lowest quartile of interbirth intervals separating live births in this population (5)), and maternal loss in the infant and juvenile period (before age 4, the earliest age of maturation in the Amboseli baboons) (72).

During the 1970s and 1980s, the quality of resources in the baboons’ habitat markedly degraded leading up to a shift in home range in the early 1990s (52). We therefore also considered a binary measure of habitat quality at birth, based on the subject’s birthdate: individuals born prior to this home range shift were considered to have been born in low-quality habitat and individuals born after the home range shift were considered to have been born in high-quality habitat. Dominance rank was estimated using ordinal ranks, where the highest status animal is given a value of 1 and individuals lower in the hierarchy have progressively larger values (63). Dominance ranks in Amboseli are determined on a monthly basis from the outcomes of dyadic agonistic interactions observed in same month. For 98% of individuals, age was based on direct observation of birth events, to within a few days’ error (SI methods).

For each CpG site, we modeled variation in DNA methylation at each CpG site in our analysis set using the binomial mixed-effects model implemented in *MACAU* (58). We controlled for genetic relatedness between individuals using genotype data derived from low-coverage resequencing data generated for all individuals in our sample in previous work (81) (SI Methods). We controlled for technical effects (e.g., batch, sequencing depth, bisulfite conversion rate) as additional fixed effects and kinship/population structure using a random effect. Using a subset of our data, we also confirmed that major differences in cell composition (lymphocyte and monocyte ratios, available from blood smear data) do not significantly predict DNA methylation in our sample (SI methods). We did not model an effect of sex because in preliminary analysis, we observed little to no signature of sex in the DNA methylation data, consistent with (82). ChromHMM tracts were based on orthology to annotations in human PBMCs generated by the Roadmap Epigenomics Consortium and converted to baboon genome coordinates using *liftOver* (27, 83). Measures of regulatory activity were assayed using mSTARR-seq on baboon DNA fragments following (29) (SI Methods). Gene expression measures from leukocytes for the same population were generated previously (60, 66).

All statistical analyses in this work were performed in R (Version #4.1.2) (84), with code available at https://github.com/janderson94/Anderson_et_al_socioecological_methylation_predictors. Newly generated RRBS sequence data have been deposited in the NCBI Short Read Archive (SRA project # PRJNA970398). mSTARR data have been deposited under NCBI SRA project #PRJNA871297. SRA accessions for previously published data can be found in Table S1.

## Acknowledgements

We thank Jeanne Altmann for her foundational contributions to the Amboseli Baboon Research Project. We thank Raphael Mututua, Serah Sayialel, Kinyua Warutere, Long’ida Siodi, Vivian Oudu, and Tim Wango for their long-term contributions to observations and project management in Kenya; Tauras Vilgalys, Arielle Fogel, and members of the Tung Lab for constructive feedback on this work; Mike Yuan and Tawni Voyles for assistance in the lab; and Jake Gordon, Niki Learn, and Karl Pinc for their contributions to database design and maintenance. In Kenya, our research was approved by the Wildlife Research Training Institute (WRTI), Kenya Wildlife Service (KWS), the National Commission for Science, Technology, and Innovation (NACOSTI), and the National Environment Management Authority (NEMA). We also thank the University of Nairobi, the Institute of Primate Research (IPR), the National Museums of Kenya, the members of the Amboseli-Longido pastoralist communities, the Enduimet Wildlife Management Area, Ker & Downey Safaris, Air Kenya, and Safarilink for their cooperation and assistance in the field, and Duke University, Princeton University, the University of Notre Dame, and the Max Planck Institute for Evolutionary Anthropology for financial and logistical support. This work was supported by National Institutes of Health R01HD088558, the Canadian Institute for Advanced Research, National Science Foundation BCS-04161, the Leakey Foundation, nad high-performance computing resources supported by the North Carolina Biotechnology Center (2016-IDG-103 and 2020-IIG-2109). We also gratefully acknowledge the NSF and NIH for support for the long-term data, currently through NSF IOS 1456832, NIH R01AG053308, R01AG053330, R01AG071684, R01AG075914, and P01AG031719.

## References

1. T. Roseboom, S. de Rooij, R. Painter, The Dutch famine and its long-term consequences for adult health. Early Hum. Dev. 82, 485–491 (2006).

2. V. J. Felitti, et al., Relationship of childhood abuse and household dysfunction to many of the leading causes of death in adults: The Adverse Childhood Experiences (ACE) Study. Am. J. Prev. Med. 14, 245–258 (1998).

3. D. W. Brown, et al., Adverse childhood experiences and the risk of premature mortality. Am. J. Prev. Med. 37, 389–396 (2009).

4. E. D. Strauss, D. Shizuka, K. E. Holekamp, Juvenile rank acquisition is associated with fitness independent of adult rank. Proc. R. Soc. B 287, 20192969 (2020).

5. J. Tung, E. A. Archie, J. Altmann, S. C. Alberts, Cumulative early life adversity predicts longevity in wild baboons. Nat. Commun. 7, 1–7 (2016).

6. G. Pigeon, F. Pelletier, Direct and indirect effects of early-life environment on lifetime fitness of bighorn ewes. Proc. R. Soc. B Biol. Sci. 285, 20171935 (2018).

7. C. R. Pryce, et al., Long-term effects of early-life environmental manipulations in rodents and primates: potential animal models in depression research. Neurosci. Biobehav. Rev. 29, 649–674 (2005).

8. D. Kaufman, et al., Early-life stress and the development of obesity and insulin resistance in juvenile bonnet macaques. Diabetes 56, 1382–1386 (2007).

9. K. L. Brunson, et al., Mechanisms of late-onset cognitive decline after early-life stress. J. Neurosci. 25, 9328–9338 (2005).

10. M. M. Sanchez, C. O. Ladd, P. M. Plotsky, Early adverse experience as a developmental risk factor for later psychopathology: evidence from rodent and primate models. Dev. Psychopathol. 13, 419–449 (2001).

11. A. M. Dettmer, J. J. Heckman, J. Pantano, V. Ronda, S. J. Suomi, “Intergenerational Effects of Early-Life Advantage: Lessons from a Primate Study” (National Bureau of Economic Research, 2020).

12. G. Conti, et al., Primate evidence on the late health effects of early-life adversity. Proc. Natl. Acad. Sci. 109, 8866–8871 (2012).

13. G. E. Miller, et al., Low early-life social class leaves a biological residue manifested by decreased glucocorticoid and increased proinflammatory signaling. Proc. Natl. Acad. Sci. 106, 14716–14721 (2009).

14. M. M. Kittleson, et al., Association of childhood socioeconomic status with subsequent coronary heart disease in physicians. Arch. Intern. Med. 166, 2356–2361 (2006).

15. C. Hertzman, Putting the concept of biological embedding in historical perspective. Proc. Natl. Acad. Sci. 109, 17160–17167 (2012).

16. A. E. Berens, S. K. G. Jensen, C. A. Nelson, Biological embedding of childhood adversity: from physiological mechanisms to clinical implications. BMC Med. 15, 1–12 (2017).

17. C. A. Demetriou, et al., Biological embedding of early-life exposures and disease risk in humans: a role for DNA methylation. Eur. J. Clin. Invest. 45, 303–332 (2015).

18. A. Bird, DNA methylation patterns and epigenetic memory. Genes Dev. 16, 6–21 (2002).

19. M. Szyf, The early life social environment and DNA methylation: DNA methylation mediating the long-term impact of social environments early in life. Epigenetics 6, 971–978 (2011).

20. M. Szyf, The early-life social environment and DNA methylation. Clin. Genet. 81, 341–349 (2012).

21. A. Pacis, et al., Gene activation precedes DNA demethylation in response to infection in human dendritic cells. Proc. Natl. Acad. Sci. 116, 6938 LP –6943 (2019).

22. S. Sun, L. B. Barreiro, The epigenetically-encoded memory of the innate immune system. Curr. Opin. Immunol. 65, 7–13 (2020).

23. S. Voisin, N. Eynon, X. Yan, D. J. Bishop, Exercise training and DNA methylation in humans. Acta Physiol. 213, 39–59 (2015).

24. N. Provençal, et al., Glucocorticoid exposure during hippocampal neurogenesis primes future stress response by inducing changes in DNA methylation. Proc. Natl. Acad. Sci. 117, 23280–23285 (2020).

25. A. Q. Fu, D. P. Genereux, R. Stöger, C. D. Laird, M. Stephens, Statistical inference of transmission fidelity of DNA methylation patterns over somatic cell divisions in mammals. Ann. Appl. Stat. 4, 871 (2010).

26. Z. Siegfried, I. Simon, DNA methylation and gene expression. Wiley Interdiscip. Rev. Syst. Biol. Med. 2, 362–371 (2010).

27. A. Kundaje, et al., Integrative analysis of 111 reference human epigenomes. Nature 518, 317–330 (2015).

28. A. Vojta, et al., Repurposing the CRISPR-Cas9 system for targeted DNA methylation. Nucleic Acids Res. 44, 5615–5628 (2016).

29. A. J. Lea, et al., Genome-wide quantification of the effects of DNA methylation on human gene regulation. Elife 7, e37513 (2018).

30. M. L. Maeder, et al., Targeted DNA demethylation and activation of endogenous genes using programmable TALE-TET1 fusion proteins. Nat. Biotechnol. 31, 1137–1142 (2013).

31. E. Kreibich, R. Kleinendorst, G. Barzaghi, S. Kaspar, A. R. Krebs, Single-molecule footprinting identifies context-dependent regulation of enhancers by DNA methylation. Mol. Cell 83, 787–802 (2023).

32. J. C. Klein, A. Keith, V. Agarwal, T. Durham, J. Shendure, Functional characterization of enhancer evolution in the primate lineage. Genome Biol. 19, 99 (2018).

33. C. V Breton, et al., Small-magnitude effect sizes in epigenetic end points are important in children’s environmental health studies: the children’s environmental health and disease prevention research center’s epigenetics working group. Environ. Health Perspect. 125, 511–526 (2017).

34. E. C. Dunn, et al., Sensitive periods for the effect of childhood adversity on DNA methylation: results from a prospective, longitudinal study. Biol. Psychiatry 85, 838–849 (2019).

35. L. C. Houtepen, et al., Childhood adversity and DNA methylation in two population-based cohorts. Transl. Psychiatry 8, 1–12 (2018).

36. C. A. M. Cecil, Y. Zhang, T. Nolte, Childhood maltreatment and DNA methylation: a systematic review. Neurosci. Biobehav. Rev. 112, 392–409 (2020).

37. Z. Stein, M. Susser, G. Saenger, F. Marolla, Famine and human development: The Dutch hunger winter of 1944-1945. (1975).

38. E. W. Tobi, et al., DNA methylation signatures link prenatal famine exposure to growth and metabolism. Nat. Commun. 5, 1–14 (2014).

39. R. A. Waterland, et al., Season of conception in rural gambia affects DNA methylation at putative human metastable epialleles. PLoS Genet. 6, e1001252 (2010).

40. A. A. Lussier, et al., Sensitive periods for the effect of childhood adversity on DNA methylation: Updated results from a prospective, longitudinal study. Biol. Psychiatry Glob. Open Sci. (2022).

41. Z. M. Laubach, et al., Early life social and ecological determinants of global DNA methylation in wild spotted hyenas. Mol. Ecol. 28, 3799–3812 (2019).

42. Z. M. Laubach, et al., Early-life social experience affects offspring DNA methylation and later life stress phenotype. Nat. Commun. 12, 1–15 (2021).

43. C. Catale, et al., Exposure to different early-life stress experiences results in differentially altered DNA methylation in the brain and immune system. Neurobiol. Stress 13, 100249 (2020).

44. S. C. Alberts, J. Altmann, “The Amboseli Baboon Research Project: 40 years of continuity and change” in Long-Term Field Studies of Primates, (Springer, 2012), pp. 261–287.

45. M. N. Zipple, E. A. Archie, J. Tung, J. Altmann, S. C. Alberts, Intergenerational effects of early adversity on survival in wild baboons. Elife 8 (2019).

46. A. J. Lea, J. Altmann, S. C. Alberts, J. Tung, Developmental Constraints in a Wild Primate. Am. Nat. 185, 809–821 (2015).

47. S. K. Patterson, S. C. Strum, J. B. Silk, Early life adversity has long-term effects on sociality and interaction style in female baboons. Proc. R. Soc. B 289, 20212244 (2022).

48. S. K. Patterson, et al., Effects of early life adversity on maternal effort and glucocorticoids in wild olive baboons. Behav. Ecol. Sociobiol. 75, 1–18 (2021).

49. M. N. Zipple, et al., Maternal death and offspring fitness in multiple wild primates. Proc. Natl. Acad. Sci. 118 (2021).

50. S. Rosenbaum, et al., Social bonds do not mediate the relationship between early adversity and adult glucocorticoids in wild baboons. Proc. Natl. Acad. Sci. 117, 20052–20062 (2020).

51. C. J. Weibel, J. Tung, S. C. Alberts, E. A. Archie, Accelerated reproduction is not an adaptive response to early-life adversity in wild baboons. Proc. Natl. Acad. Sci. 117, 24909–24919 (2020).

52. L. R. Gesquiere, J. Altmann, E. A. Archie, S. C. Alberts, Interbirth intervals in wild baboons: Environmental predictors and hormonal correlates. Am. J. Phys. Anthropol. 166, 107–126 (2018).

53. J. Altmann, S. C. Alberts, Variability in reproductive success viewed from a life-history perspective in baboons. Am. J. Hum. Biol. 15, 401–409 (2003).

54. J. A. Anderson, et al., Distinct gene regulatory signatures of dominance rank and social bond strength in wild baboons. bioRxiv, 2021.05.31.446340 (2021).

55. N. Snyder-Mackler, et al., Social status alters immune regulation and response to infection in macaques. Science (80-.). 354, 1041 LP –1045 (2016).

56. N. Snyder-Mackler, et al., Social status alters chromatin accessibility and the gene regulatory response to glucocorticoid stimulation in rhesus macaques. Proc. Natl. Acad. Sci. U. S. A. 116, 1219–1228 (2019).

57. J. Sanz, et al., Social history and exposure to pathogen signals modulate social status effects on gene regulation in rhesus macaques. Proc. Natl. Acad. Sci., 201820846 (2019).

58. A. J. Lea, J. Tung, X. Zhou, A Flexible, Efficient Binomial Mixed Model for Identifying Differential DNA Methylation in Bisulfite Sequencing Data. PLoS Genet. 11, 1–31 (2015).

59. M. Jung, G. P. Pfeifer, Aging and DNA methylation. BMC Biol. 13, 1–8 (2015).

60. J. A. Anderson, et al., Distinct gene regulatory signatures of dominance rank and social bond strength in wild baboons. Philos. Trans. R. Soc. B 377, 20200441 (2022).

61. A. Meissner, et al., Genome-scale DNA methylation maps of pluripotent and differentiated cells. Nature 454, 766–770 (2008).

62. P. Boyle, et al., Gel-free multiplexed reduced representation bisulfite sequencing for large-scale DNA methylation profiling. Genome Biol. 13, 1–10 (2012).

63. E. J. Levy, et al., A comparison of dominance rank metrics reveals multiple competitive landscapes in an animal society. Proc. R. Soc. B 287, 20201013 (2020).

64. S. C. Alberts, H. E. Watts, J. Altmann, Queuing and queue-jumping: long-term patterns of reproductive skew in male savannah baboons, Papio cynocephalus. Anim. Behav. 65, 821–840 (2003).

65. A. J. Lea, N. H. Learn, M. J. Theus, J. Altmann, S. C. Alberts, Complex sources of variance in female dominance rank in a nepotistic society. Anim. Behav. 94, 87–99 (2014).

66. A. J. Lea, et al., Dominance rank-associated gene expression is widespread, sex-specific, and a precursor to high social status in wild male baboons. Proc. Natl. Acad. Sci. 115, E12163–E12171 (2018).

67. J. A. Anderson, et al., High social status males experience accelerated epigenetic aging in wild baboons. Elife 10, e66128 (2021).

68. Å. Johansson, S. Enroth, U. Gyllensten, Continuous aging of the human DNA methylome throughout the human lifespan. PLoS One 8, e67378 (2013).

69. S. S. Batra, et al., Accurate assembly of the olive baboon (Papio anubis) genome using long-read and Hi-C data. bioRxiv, 678771 (2019).

70. J. Friedman, T. Hastie, R. Tibshirani, Regularization paths for generalized linear models via coordinate descent. J. Stat. Softw. 33, 1 (2010).

71. K. A. McLaughlin, M. A. Sheridan, H. K. Lambert, Childhood adversity and neural development: deprivation and threat as distinct dimensions of early experience. Neurosci. Biobehav. Rev. 47, 578–591 (2014).

72. M. J. E. Charpentier, J. Tung, J. Altmann, S. C. Alberts, Age at maturity in wild baboons: Genetic, environmental and demographic influences. Mol. Ecol. 17, 2026–2040 (2008).

73. E. W. Tobi, et al., DNA methylation as a mediator of the association between prenatal adversity and risk factors for metabolic disease in adulthood. Sci. Adv. 4, eaao4364 (2018).

74. E. C. Briggs, L. Amaya-Jackson, K. T. Putnam, F. W. Putnam, All adverse childhood experiences are not equal: The contribution of synergy to adverse childhood experience scores. Am. Psychol. 76, 243 (2021).

75. Y. Yin, et al., Impact of cytosine methylation on DNA binding specificities of human transcription factors. Science (80-.). 356, eaaj2239 (2017).

76. E. C. Lange, et al., Early life adversity and adult social relationships have independent effects on survival in a wild animal model of aging. bioRxiv, 2009–2022 (2022).

77. A. F. McRae, et al., Contribution of genetic variation to transgenerational inheritance of DNA methylation. Genome Biol. 15, 1–10 (2014).

78. E. Elliott, G. Ezra-Nevo, L. Regev, A. Neufeld-Cohen, A. Chen, Resilience to social stress coincides with functional DNA methylation of the Crf gene in adult mice. Nat. Neurosci. 13, 1351–1353 (2010).

79. J. Tung, X. Zhou, S. C. Alberts, M. Stephens, Y. Gilad, The genetic architecture of gene expression levels in wild baboons. Elife 4, e04729.%@ 2050-084X (2015).

80. T. P. Vilgalys, et al., Selection against admixture and gene regulatory divergence in a long-term primate field study. Science (80-.). 377, 635–641 (2022).

81. T. P. Vilgalys, et al., Selection against admixture and gene regulatory divergence in a long-term primate field study. bioRxiv (2021).

82. A. J. Lea, J. Altmann, S. C. Alberts, J. Tung, Resource base influences genome-wide DNA methylation levels in wild baboons (Papio cynocephalus). Mol. Ecol. 25, 1681–1696 (2016).

83. A. S. Hinrichs, et al., The UCSC genome browser database: update 2006. Nucleic Acids Res. 34, D590–D598 (2006).

84. R Team, R: A language and environment for statistical computing (2013).

