## Supplementary Information for "DNA methylation signatures of early life adversity are exposure-dependent in wild baboons"

**Table of Contents**


1. Supplementary Methods
    - *DNA methylation data generation and pre-processing*
    - *Demographic, social, and ecological variables*
    - *Modeling socioenvironmental predictors of DNA methylation*
    - *Genome annotations*
    - *Elastic net regularization models*
    - *mSTARR-seq experiment*
    - *mSTARR-seq data analysis*
    - *Comparing DNA methylation to gene expression*
    - *Assessing the effects of cell-type heterogeneity*

2. Supplementary Figures
    - Figure S1. Distribution of births and sampling over time with respect to habitat shifts
    - Figure S2. Pairwise correlation of early life variables across individuals
    - Figure S3. Rainfall in the first year of life and in the year leading up to darting are weakly correlated
    - Figure S4. Overlap between age and rank effects and the effects of early life environment
    - Figure S5. Dominance rank associations with DNA methylation predict dominance rank associations with gene expression in the Amboseli baboons

3. Supplementary Tables
    - Table S1. Metadata for the DNA methylation dataset.
    - Table S2a. MACAU results for model 1.
    - Table S2b. MACAU results for model 2.
    - Table S2c. MACAU results for model 3.
    - Table S3. Site-specific enrichment of socioecological effects in various genomic compartments.
    - Table S4. Metadata for the mSTARR dataset.
    - Table S5. mSTARR model results for regulatory activity and methylation-dependent activity in tested genomic windows.

- Table S6. Enrichment of mSTARR regulatory windows in chromHMM compartments.
- Table S7. Gene Set Enrichment Analysis results of male rank effects on DNA methylation.

4. Supplementary References

### Supplementary Methods

*DNA methylation data generation and pre-processing*

DNA was extracted from whole blood using Qiagen DNeasy Blood & Tissue extraction kits following the manufacturer's instructions. Previously published data (N=264 samples) were generated using reduced representation bisulfite sequencing (RRBS) based on the protocols of (1, 2). Libraries were generated from 200 ng of DNA per sample followed by high-throughput sequencing on the Illumina HiSeq2500 or HiSeq4000 platform. Data newly generated for this study (N=31 samples) were produced using a modification of the standard RRBS protocol, double-digest RRBS (following (3)), and sequenced on the Illumina HiSeq 2500. Whereas RRBS is based on a single-step digest using the restriction enzyme *MspI*, dRRBS uses a double restriction enzyme digest (here, *MspI* and *ApeKI*) to enrich the resulting library for CpG sites outside of promoters and CpG islands. The batch used to generate the RRBS libraries and sequencing data, which also controls for differences between the dRRBS versus RRBS preps, was therefore included as a covariate in our analyses (n=7 batches; Table S1).

Both RRBS and dRRBS libraries were prepared using 0.2 ng of a lambda phage DNA spike-in, which allowed us to estimate bisulfite conversion efficiency based on reads mapped to the lambda phage genome (mean bisulfite conversion rate=0.998 $\pm$ $2.4 \times 10^{-3}$ s.d.; Table S1). Raw reads from all samples were trimmed for Illumina adaptors using TrimGalore (length=15, stringency=4) (4), and mapped to the *Panubis1.0* genome (GCA_008728515.1) using BSMAP (10% max mismatch, unique hits) (5, 6). For further analysis, we retained CpG sites (i) with non-zero coverage in at least 75% of samples; and (ii) median coverage >5 across all samples. We also excluded sites that were constitutively hypomethylated (mean methylation ratio <0.1) or constitutively hypermethylated (mean methylation ratio >0.9) in the data set, resulting in a final analysis set of 477,270 sites.

*Demographic, social, and ecological variables*

*Age* is known to within a few days' error for 251 (98%) of our study subjects, and within 6 months' error for the remaining five individuals in the data set (2% of unique individuals). Age information is based on longitudinal observations of births within study groups.

*Dominance rank* is estimated using ordinal ranks (where 1 indicates the highest status individual and progressively higher numbers correspond to progressively lower status). Males and females are ordered in distinct linear dominance hierarchies, so male rank and female rank were modeled as separate effects. Dominance ranks are based on observations of agonistic wins and losses recorded during representative interaction sampling (7, 8). In this approach, agonisms are recorded for all study group members during the course of random-order focal sampling: that is, observers collect agonism data for all animals in their line of sight while moving through the group to find and follow predetermined focal sampling subjects.

Ranks are assigned by generating an N x N matrix, where N is the number of individuals in the social group. The matrix contains symmetrical rows and columns corresponding to individual animal identities. The cells of the matrix contain the number of times that the animal represented in a given row won an agonistic interaction against the animal represented in a given column during a month-long period of data collection. The columns and rows of the matrix are then ordered to minimize the number of wins that appear below the diagonal of the matrix. The resulting order of the columns is the ordinal rank (e.g. 1, 2, 3, etc.) of the animals represented by those columns. We modeled dominance rank in adulthood based on assigned ordinal ranks in the month in which blood samples were collected, and maternal social status based on the focal animal's mother's dominance rank in the month that animal was born. For a detailed treatment of rank assignment, please see (8).

*Habitat quality* was defined as low quality before the home range shifts and high quality after the home range shifts. The two social groups that were observed during the period of the home range shifts (Alto's and Hook's groups) made the shift in different years. Hence, for Alto's group, we coded low-quality habitat based on a birthdate during or before 1987, but coded low-quality habitat for animals in Hook's group based on a birthdate during or before 1991. Animals in all other social groups in this study were born post-range shift, in high-quality habitat.

*Early life adversity.* To quantify five, individually variable dimensions of early life adversity, we followed previous studies of early adversity in the Amboseli baboons (9, 10). Specifically, we considered our study subjects to be exposed to (i) *drought* if they experienced <200 mm of rainfall in the first year of life; (ii) *maternal loss* if they lost their mother prior to 4 years of age (the earliest age of reproductive maturation in our study population); (iii) *low early life social status*, if their mother's rank at birth fell in the lowest quartile of ordinal dominance rank values (rank ≥12); (iv) a *close-in-age younger sibling* if they experienced the birth of a live younger sibling within 1.5 years of their own birth (i.e., the lowest quartile of interbirth intervals in this population); and (v) *large group size*, a measure of resource competition, if the number of adult baboons residing in their social group was in the top quartile of group size values for this population (group size ≥36). *Cumulative early adversity* was defined as the sum of exposures to these individual sources of adversity and ranged from 0 to 4 in our sample (median=1; s.d.=0.97). Note that in this analysis, we omitted a sixth source of early adversity, maternal social isolation, which was included in (9). This measure is most prone to missingness in the data set, so we followed the precedent in (10, 11), which maximizes the analysis set using a five-exposure cumulative early adversity index.

*Modeling socioenvironmental predictors of DNA methylation*

We modeled variation in DNA methylation at each CpG site in our analysis set using the binomial mixed-effects model implemented in *MACAU*, which is designed specifically for bisulfite sequencing data (12). The basic form of the model for each CpG site is:

$$y_i \sim Bin(r_i, \pi_i)$$

where $r_i$ is the total read count for individual *i*, $y_i$ is the methylated read count, and $\pi_i$ is the true, unknown underlying proportion of methylated reads for individual *i*. $\pi_i$ is passed through a logit link and modelled as:

$$Logit\left(\frac{\pi_i}{1-\pi_i}\right) = w_i^T \alpha + x_i^T \beta + g_i + e_i$$

$$g \sim MVN(0, \ \sigma^2 h^2 K)$$
$$e \sim MVN(0, \ \sigma^2 (1-h^2) I)$$

where: w is an n x m-matrix of covariates, including an intercept; α is the corresponding m-vector of coefficients; $x_i$ is a n by p-matrix of predictors of interest for individual *i*; $\beta$ is the corresponding p-vector of coefficients; $g_i$ is an n-vector of random effect estimates that capture the effect of kinship or shared ancestry; MVN is the multivariate normal distribution; $\sigma^2 h^2$ is the genetic variance component; *K* is a genetic relatedness matrix; *e* is an n-vector of residual errors; $\sigma^2(1-h^2)$ is the environmental variance component; and *I* is the identity matrix. Note that the *w* and *x* vectors are both modeled as fixed effects. We separate them here conceptually to distinguish between variables whose effects we are interested in controlling for (*w*), and those we are directly interested in estimating and interpreting (*x*).

We fit three related models to our data. All three models used the same random effects structure and incorporated the same *w* matrix, including the technical effects of z-scored bisulfite conversion rate, z-scored sequencing depth, and sampling batch. Sampling batch assignment was based on the batch in which the sample library was generated and sequenced (n=7 batches, which also capture the differences in dRRBS/RRBS library preparation). To estimate the genetic relatedness matrix *K*, we calculated the variance-covariance matrix of genotype data for the individuals in our sample, rescaled so that the trace(*K*)=1 (12). Genotype data were derived from low-coverage resequencing data generated for all individuals in our sample in previous work (See SI section 4 in (13)). In brief, variants were jointly genotyped using the Genome Analysis Toolkit (14), after removing PCR duplicates. Low quality genotypes were removed, filtered for minor allele frequency >0.05, and thinned by 100,000 base pairs, resulting in 25,628 biallelic SNPs. We defined the K matrix for our analyses as the variance-covariance matrix of the genotypes at these loci. Each model differed only in the composition of the matrix *x*. Thus our three models took the following forms:

$$Model\ 1: Logit\left(\frac{\pi_i}{1-\pi_i}\right) = w_i^T\alpha + x_{model1\_i}^T\beta + g_i + e_i$$

$$Model\ 2: Logit\left(\frac{\pi_i}{1-\pi_i}\right) = w_i^T\alpha + x_{model2\_i}^T\beta + g_i + e_i$$

$$Model\ 3: Logit\left(\frac{\pi_i}{1-\pi_i}\right) = w_i^T\alpha + x_{model3\_i}^T\beta + g_i + e_i$$

where,

$$x_{model1} = Age, HQ\ , Rank\ (sex = M), Rank(sex = F),\ CEA$$
$$x_{model2} = Age, HQ\ , CEA\ (HQ = 0),\ CEA\ (HQ = 1)$$
$$x_{model3} = Age, HQ, [EA_1(HQ = 0), EA_1(HQ = 1)] \ldots [EA_5(HQ = 0), EA_5(HQ = 1)]$$

and *Age* is a continuous measure of age in years; *HQ* is a binary 0/1 variable capturing early life habitat quality (0=high quality habitat; 1=low quality habitat); *CEA* is cumulative early adversity, represented as an integer value from 0 to 5; *Rank* is an individual's sex-specific ordinal dominance rank at the time of sampling; and $EA_n$ represents a series of binary variables that reflect an individual's exposure to each of five forms of early adversity (maternal loss, low maternal social status, a close-in-age younger sibling, high experience density/group size, drought in the first year of life). Age and sex-specific dominance rank values were z-scored across samples prior to modeling. In all cases, we tested the hypothesis that the effect size for each variable of interest did not equal zero. To control for multiple hypothesis testing, we used the false discovery rate approach implemented in the R package *qvalue* (15, 16), after confirming that permutations of our predictors of interest generated null p-value distributions similar to the uniform distribution.

*Genome annotations*

Gene bodies were defined based on annotations for the baboon genome (*Panubis1.0* GTF # GCF_008728515.1) (6). Promoters were defined as the 2 kb upstream of a gene's 5'-most annotated transcription start site. CpG islands were defined as windows longer than 200 bp with greater than 50% GC content and an observed/expected CpG ratio greater than 0.6, as identified using *EMBOSS* (17). CpG shores were annotated based on the 2 kb regions upstream and downstream of CpG islands. Finally, putative enhancer elements were identified based on *liftOver* (18, 19) of H3K4me1 ChIP-seq peaks from human PBMCs, generated by the ENCODE project (experiment ENCSR482QXO) (20).

To define chromatin states, we used chromatin state annotations in human peripheral blood mononuclear cells generated using chromHMM (21), which are based on five histone marks (H3K4me3, H3K4me1, H3K36me3, H3K27me3, H3K9me3). As for H3K4me1-defined enhancers, we used *liftOver* to identify regions in the baboon genome that correspond to calls in humans, based on 200 bp non-overlapping windows of the human genome (18). In both cases, we used default *liftOver* parameters, and only retained regions that resulted in unique hits when reciprocally lifting over from the human genome to the baboon genome as well as back to the same human coordinates (19). The liftover chainfile generated previously is available at https://zenodo.org/record/5199534#.Y_FamezML0p.

*Elastic net regularization models*

We predicted early life habitat quality status (i.e. pre- versus post-shift) for each sample using elastic net regression in the R package *glmnet* (22). Specifically, we imputed missing methylation ratios (<5%) for each sample using the R package *impute* (23). We then iteratively removed one sample at a time and trained an elastic net model on the remaining training set using 50-fold internal cross-validation, an alpha value of 1, and the lambda value that minimized mean-squared error during internal CV. The resulting model was then used to predict habitat quality (low versus high) for the originally removed test set sample. We repeated this process for each sample to obtain an estimate of accuracy and an ROC curve.

To test whether predictive ability declines with time since the shift from low to high-quality habitat, we used a linear model to model predicted habitat quality as a function of the time between sample collection and when the animal left their low-quality habitat. Because of the natural correlation between the time since habitat shift and animals' ages, we cannot effectively control for the animals' ages. We note, however, that predicted habitat quality is not significantly associated with age in a linear model. Thus, our model is unlikely to be capturing an age effect rather than difference in habitat quality.

*mSTARR-seq experiment*

To prepare plasmid libraries for mSTARR-seq transfection, we extracted genomic DNA from cryopreserved peripheral blood mononuclear cells (Qiagen, Blood and Cell Culture DNA Mini Kit). The cells were sampled from individual #15944 of the Southwest National Primate Research Center, the same anubis baboon that was used to generate the *Panubis1.0* genome assembly (6). 300-800 bp DNA fragments were generated in two ways: (i) using a Covaris S220 Focused-Ultrasonicator followed by size selection, which represents fragments sampled from across the entire genome ("sheared library"); and (ii) via digestion with the restriction enzyme *Msp1*, also followed by size selection, which generates fragments that are enriched in baboon RRBS libraries ("*MspI*-digested library"). The *MspI*-digested library mimics the first step of the RRBS protocol, which also involves *MspI* digestion. By generating both types of libraries, our goal was to enrich for fragments that we measured in the Amboseli baboon data set while also capturing fragments representative of the genome as a whole.

Plasmid libraries, transfection, and harvest protocols followed the published protocol from (24). In brief, size-selected fragments were ligated to NEBNext adapters (NEB #E7335), amplified with primers complementary to the insert sites for *pmSTARRseq1* (the CpG-free plasmid backbone used for mSTARR-seq assays), and cloned into the *pmSTARRseq1* backbone using Gibson assembly. We then transformed the libraries into customized electrocompetent GT115 *E. coli* cells (300 μl, Intact Genomics), incubated them overnight at 37°C, and purified the plasmid pool (Qiagen Plasmid Plus Maxi Kit). We initially performed 10 replicate transformations each for the sheared and *MspI*-digested libraries. After estimating fragment diversity in each replicate via sequencing on an Illumina MiSeq (paired-end 75 bp reads; Table S4), we constructed our final libraries by pooling 300 ug each of the two most diverse *MspI*-digested replicates into an "*MspI*" pool and 120 ug each of the five most diverse sheared library replicates into a "sheared" pool.

To create matched unmethylated and methylated libraries, we split each pool in half and treated one half with 150U of the enzyme *M.SssI* (New England Biolabs), which methylates all CpG sites on the fragment inserts (the backbone is CpG free) and the other half with water, which leaves all CpG sites in the inserts unmethylated. Methylated versions of the Msp1-digested and sheared libraries were mixed in a 1:1 ratio, and unmethylated versions of the Msp1-digested and sheared libraries were also mixed 1:1. Following the published mSTARR-seq protocol (18), we then performed chemical transfection (Thermo Fisher Scientific Lipofectamine 3000) of 40 ug of either the methylated or unmethylated plasmid libraries into the human K562 erythroleukemic cell line, in six replicates per treatment (ca. 20 million cells). After a 48 hour incubation in opti-MEM culture media, we harvested the cells and used a quarter of the final cell suspension (in PBS) to purify plasmids for DNA-seq to quantify input for each region and the rest for mSTARR-seq plasmid-specific RNA-seq to measure each region's enhancer-like activity. Both DNA-seq and RNA-seq libraries were specifically targeted to fragment inserts and transcripts produced from the plasmid, respectively, using targeted PCR and the KAPA HiFi HotStart ReadyMix (Roche) (24). DNA-seq (n=6 replicates each from the methylated and unmethylated treatments) and RNA-seq (n=6 replicates each from the methylated and unmethylated treatments) libraries were sequenced on a NovaSeq 6000 S1 flow cell using 100 bp paired end reads. The average sequencing depth for DNA-seq libraries was 75,179,499 ± 23,113,359 reads (mean ± s.d.), and for RNA-seq libraries, the average depth was 51,520,043 ± 5,912,789 reads (mean ± s.d.) (Table S4).

*mSTARR-seq data analysis*

Raw reads were trimmed with Cutadapt (25) and Trim Galore (4) and mapped to the anubis baboon reference genome (*Panubis 1.0*) with *bwa* (*bwa mem* with default parameters) (26). We retained properly paired reads with MAPQ ≥10. Fragments that derived from the *MspI*-digested libraries versus sheared libraries were identified based on the presence of an *Msp1* cut site at the start of either the forward or reverse read. For each replicate (n=6 unmethylated DNA; n=6 methylated DNA; n=6 methylated RNA; n=6 unmethylated RNA), and separately for *MspI*-derived fragments and sheared fragments, we used *bedtools2* (27) to count the number of reads that overlapped discrete 500 bp windows in the baboon genome. We chose to use 500 bp windows for this analysis, as opposed to the original 200 bp windows in (24), because 500 bp windows maximized enrichment of ENCODE-annotated enhancer elements (lifted over to the baboon genome) among putative regulatory elements called from the experiment. Larger window sizes also reduced cases of pseudoreplication, in which we called multiple regulatory elements directly adjacent to one another, which probably function biologically as a single element.

For downstream analysis, we retained only those windows with (i) median coverage ≥ 4x in both methylated treatment DNA samples and unmethylated treatment DNA samples (i.e., where there was sufficient fragment input to drive gene expression, if capable of doing so); (ii) non-zero counts in at least half of DNA-seq replicates in *both* treatments; and (iii) non-zero counts in at least half of RNA-seq replicates in *either* treatment. The stricter criteria for DNA-seq reads is because DNA fragments must be successfully introduced into the cells to even be tested for regulatory activity. In contrast, low or no RNA-seq reads in one treatment condition, if the plasmids containing the matching DNA fragments are present, is a biological signal of the lack of regulatory potential. Following filtering, we retained 210,942 analyzable windows for the *MspI*-digested libraries and 41,521 windows for the sheared libraries, representing ~126 Mb of the baboon genome (~4% of the genome). Before testing the regulatory capability of analyzable windows, we normalized library size for each sample with *calNormFactors* function as implemented in *edgeR* package (28–30), and normalized each RNA-seq sample against its corresponding DNA-seq samples with the *voomWithQualityWeights* function implemented in the *limma* R package (31–33), so that we could later model RNA abundance relative to DNA abundance as described below.

To test for regulatory capacity and methylation-dependent regulatory activity, we fit the following model to each analyzable window:

$$y_i = \mu + m_i\beta_1 + t_i\beta_2 * I(m = 0) + t_i\beta_3 * I(m = 1) + \varepsilon_i$$

where $y_i$ is the vector of normalized counts per 500-bp window for a total of 24 samples ($n_{DNA}$=12, $n_{RNA}$=12), indexed by *i*; $\mu$ is the intercept; *m* is treatment (0=unmethylated; 1=methylated) and $\beta_1$ is its effect size; *t* is sample type (0=DNA; 1=RNA) and *I* is an indicator variable for whether the sample was unmethylated (m=0) or methylated (m=1); $\beta_2$ and $\beta_3$ are the effect sizes for sample type (RNA versus DNA) in the unmethylated and methylated conditions, respectively. $\varepsilon_i$ is the residual error. The regulatory activity for fragments produced via sheared and *MspI*-digested libraries were modeled separately. Due to the typically higher coverage in regions covered by *MspI*-digested fragments, we used results from the *MspI* digestion if coverage was available from both MspI and sheared libraries.

Regions capable of regulatory activity generate more RNA than expected based on the amount of DNA input for that region. We therefore were specifically interested in regions with positive effect sizes for the sample type (RNA versus DNA) effect, such that mRNA abundance is significantly greater than input DNA abundance for the same fragment, either in the methylated condition, unmethylated condition, or both. To control for multiple hypothesis testing, we used a permutation-based false discovery rate approach. Specifically, we randomized the DNA versus RNA label within replicate pairs, reran the model described above, and retained the same number of regions with positive RNA versus DNA effect sizes as detected in the empirical sample. We then compared the p-value distribution for these regions, across 100 permutations, to the p-values for positive effect sizes identified in the real data, using a 10% FDR cut-off (i.e. q-value <0.1) for significance (Table S5). As in previous studies, regions with significant regulatory activity detected at this threshold were enriched in strong enhancer and active promoter chromatin states annotated in K562 cells ($\log_2$(OR)=2.50 and 0.92 respectively, both $p<1 \times 10^{-9}$; Table S6), indicating that the mSTARR-seq-annotated regulatory elements are consistent with *in vivo* expectations.

Finally, to identify methylation-dependent regulatory elements, we focused on the subset of windows with regulatory activity (n=5,878 detected at q-value <0.1; Table S5). For these windows, we tested whether the effect of sample type (RNA versus DNA) differed between methylated and unmethylated conditions (i.e., whether a fragment's capacity to drive regulatory activity differs depending on whether it was methylated or not, such that $\beta_2$ and $\beta_3$ significantly differ). To correct for multiple testing in this analysis, we calculated q-values by comparing p-values from the empirical results against results from 100 permutations where treatment condition (methylated versus unmethylated) was randomly assigned to each DNA-RNA replicate pair (Table S5).

*Comparing DNA methylation to gene expression*

Male dominance rank effects on gene expression were estimated in previously published work (34). In brief, RNA-seq data were collected from white blood cells purified from *ex vivo*-incubated TruCulture tubes (Myriad RBM). Because the original study was interested in assessing sources of variance in the immune response, two TruCulture tubes were collected from each study subject: one containing cell culture media only (the “baseline” control condition) and one containing cell culture media plus lipopolysaccharide, to mimic bacterial exposure. Here, we focused on rank effect estimates in the baseline samples only (34) and on CpG sites within annotated gene bodies.

To identify pathways and gene categories more closely associated with differentially methylated CpG sites than expected by chance, we performed gene set enrichment analysis on CpG-associated genes using *GSEA v1.0* in R (35) for each of fifty Hallmark gene sets annotated in the Molecular Signatures Database (36). P-values were calculated by randomly permuting gene labels and rerunning GSEA 1000 times for each gene set. P-values for a gene set were defined by the number of permuted enrichment scores that were larger in magnitude than the empirical enrichment score.

*Assessing the effects of cell type heterogeneity*

Differential methylation can occur because of changes in methylation within cells or because of compositional effects, in which blood cell subtypes differ between, e.g., individuals exposed to high versus low early adversity, and these subtypes also differ in their DNA methylation patterns at putatively differentially methylated sites. To assess the potential confounding effects of cell-type heterogeneity in our data set, we drew on blood cell counts performed for Giemsa-stained blood smears collected in parallel with the blood samples used for DNA methylation data generation. These data, which capture the percentage of white blood cells in a sample that are monocytes, basophils, eosinophils, neutrophils, or lymphocytes, were available for 137 of our 295 samples (37). Importantly, none of these values are correlated with the major predictors of interest in our models (all $p > 0.05$ for pairwise correlations between blood cell proportions and cumulative early adversity, male rank, and drought).

We then focused on the two major cell types observed in our blood smears, neutrophils and lymphocytes. Re-running Model 1 on our data, including z-scored neutrophil and lymphocyte proportions (12), revealed few significant associations between DNA methylation and neutrophil or lymphocyte proportions (13 and 17 sites respectively < 10% FDR). Additionally, the top 5% of neutrophil and lymphocyte-associated sites do not significantly overlap with the set of habitat quality or cumulative early adversity (in low habitat quality)-associated sites identified in Model 2 (all FET p > 0.20).

### References


1. P. Boyle, *et al.*, Gel-free multiplexed reduced representation bisulfite sequencing for large-scale DNA methylation profiling. *Genome Biol.* **13**, 1–10 (2012).
2. H. Gu, *et al.*, Preparation of reduced representation bisulfite sequencing libraries for genome-scale DNA methylation profiling. *Nat. Protoc.* **6**, 468–481 (2011).
3. J. Wang, *et al.*, Double restriction-enzyme digestion improves the coverage and accuracy of genome-wide CpG methylation profiling by reduced representation bisulfite sequencing. *BMC Genomics* **14**, 1–12 (2013).
4. F. Krueger, Trim galore. *A wrapper tool around Cutadapt FastQC to consistently apply Qual. Adapt. trimming to FastQ files* **516**, 517 (2015).
5. Y. Xi, W. Li, BSMAP: whole genome bisulfite sequence MAPping program. *BMC Bioinformatics* **10**, 232 (2009).
6. S. S. Batra, *et al.*, Accurate assembly of the olive baboon (Papio anubis) genome using long-read and Hi-C data. *Gigascience* **9**, giaa134 (2020).
7. S. Alberts, *et al.*, Monitoring guide for the Amboseli Baboon Research Project (2018).
8. J. B. Gordon, D. Jansen, N. Learn, S. C. Alberts, Ordinal dominance rank assignments: Protocol for the Amboseli Baboon Research Project.
9. J. Tung, E. A. Archie, J. Altmann, S. C. Alberts, Cumulative early life adversity predicts longevity in wild baboons. *Nat. Commun.* **7**, 1–7 (2016).
10. M. N. Zipple, E. A. Archie, J. Tung, J. Altmann, S. C. Alberts, Intergenerational effects of early adversity on survival in wild baboons. *Elife* **8** (2019).
11. C. J. Weibel, J. Tung, S. C. Alberts, E. A. Archie, Accelerated reproduction is not an adaptive response to early-life adversity in wild baboons. *Proc. Natl. Acad. Sci.* **117**, 24909–24919 (2020).
12. A. J. Lea, J. Tung, X. Zhou, A Flexible, Efficient Binomial Mixed Model for Identifying Differential DNA Methylation in Bisulfite Sequencing Data. *PLoS Genet.* **11**, 1–31 (2015).
13. T. P. Vilgalys, *et al.*, Selection against admixture and gene regulatory divergence in a long-term primate field study. *Science (80-. ).* **377**, 635–641 (2022).
14. G. A. Van der Auwera, B. D. O'Connor, *Genomics in the cloud: using Docker, GATK, and WDL in Terra* (O'Reilly Media, 2020).
15. J. D. Storey, R. Tibshirani, Statistical significance for genomewide studies. *Proc. Natl. Acad. Sci.* **100**, 9440–9445 (2003).
16. A. Dabney, J. D. Storey, G. R. Warnes, qvalue: Q-value estimation for false discovery rate control. *R Packag. version* **1** (2010).
17. P. Rice, I. Longden, A. Bleasby, EMBOSS: the European molecular biology open software suite. *Trends Genet.* **16**, 276–277 (2000).
18. A. S. Hinrichs, *et al.*, The UCSC genome browser database: update 2006. *Nucleic Acids Res.* **34**, D590–D598 (2006).
19. T. P. Vilgalys, *et al.*, Selection against admixture and gene regulatory divergence in a long-term primate field study. *bioRxiv* (2021).
20. J. Zhang, *et al.*, An integrative ENCODE resource for cancer genomics. *Nat.*

*Commun.* **11**, 1–11 (2020).
21. A. Kundaje, *et al.*, Integrative analysis of 111 reference human epigenomes. *Nature* **518**, 317–330 (2015).
22. J. Friedman, T. Hastie, R. Tibshirani, Regularization paths for generalized linear models via coordinate descent. *J. Stat. Softw.* **33**, 1 (2010).
23. T. Hastie, *et al.*, Imputing missing data for gene expression arrays (1999).
24. A. J. Lea, *et al.*, Genome-wide quantification of the effects of DNA methylation on human gene regulation. *Elife* **7**, e37513 (2018).
25. M. Martin, Cutadapt removes adapter sequences from high-throughput sequencing reads. *EMBnet. J.* **17**, 10–12 (2011).
26. H. Li, Aligning sequence reads, clone sequences and assembly contigs with BWA-MEM. *arXiv Prepr. arXiv1303.3997* (2013).
27. A. R. Quinlan, I. M. Hall, BEDTools: a flexible suite of utilities for comparing genomic features. *Bioinformatics* **26**, 841–842 (2010).
28. S. Anders, W. Huber, Differential expression analysis for sequence count data. *Genome Biol* **11** (2010).
29. J. H. Bullard, E. Purdom, K. D. Hansen, S. Dudoit, Evaluation of statistical methods for normalization and differential expression in mRNA-Seq experiments. *BMC Bioinformatics* **11**, 1–13 (2010).
30. M. D. Robinson, A. Oshlack, A scaling normalization method for differential expression analysis of RNA-seq data. *Genome Biol.* **11**, 1–9 (2010).
31. C. W. Law, Y. Chen, W. Shi, G. K. Smyth, voom: Precision weights unlock linear model analysis tools for RNA-seq read counts. *Genome Biol.* **15**, R29 (2014).
32. R. Liu, *et al.*, Why weight? Modelling sample and observational level variability improves power in RNA-seq analyses. *Nucleic Acids Res.* **43**, e97–e97 (2015).
33. M. E. Ritchie, *et al.*, Empirical array quality weights in the analysis of microarray data. *BMC Bioinformatics* **7**, 1–16 (2006).
34. J. A. Anderson, *et al.*, Distinct gene regulatory signatures of dominance rank and social bond strength in wild baboons. *Philos. Trans. R. Soc. B* **377**, 20200441 (2022).
35. A. Subramanian, *et al.*, Gene set enrichment analysis: a knowledge-based approach for interpreting genome-wide expression profiles. *Proc. Natl. Acad. Sci.* **102**, 15545–15550 (2005).
36. A. Liberzon, *et al.*, The molecular signatures database hallmark gene set collection. *Cell Syst.* **1**, 417–425 (2015).
37. A. J. Lea, J. Altmann, S. C. Alberts, J. Tung, Resource base influences genome-wide DNA methylation levels in wild baboons (Papio cynocephalus). *Mol. Ecol.* **25**, 1681–1696 (2016).

**Supplementary Figures**

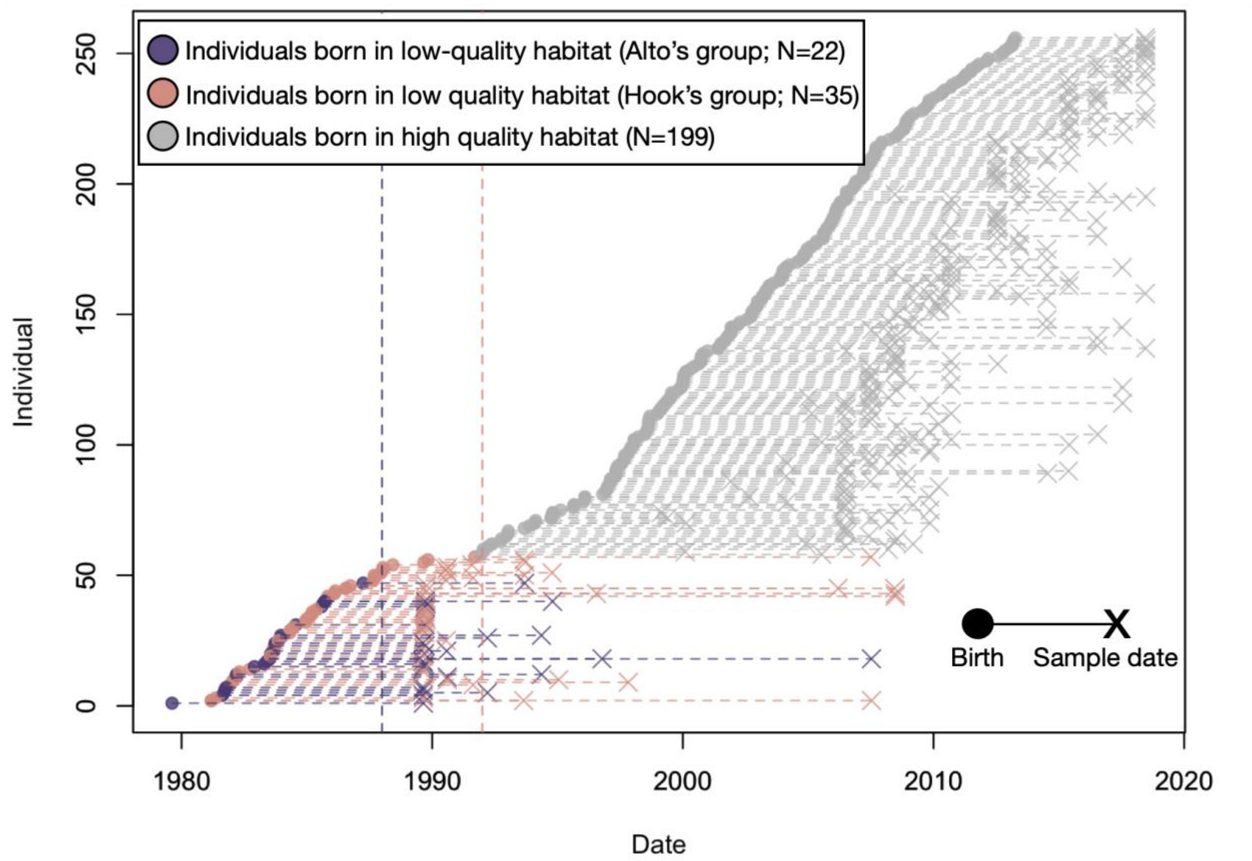


**Figure S1: Distribution of births and sampling dates with respect to habitat shifts.** Each individual is represented by a dashed horizontal line, ordered on the y-axis based on date of birth. Closed circles at the left end of each line show birth dates and x's at the right end of each line show blood sample date (in 37 cases, animals were sampled multiple times, so multiple x's occur on those lines). Colored lines show animals born in the low-quality habitat (colored dots and lines); gray lines show animals born in the high-quality habitat. The two social groups that were studied before the habitat shift (Hook's group and Alto's group) are colored in peach and purple, respectively. Vertical dashed lines show the year in which Alto's group and then Hook's group shifted from low-quality to high-quality habitat (1988 and 1992 respectively).

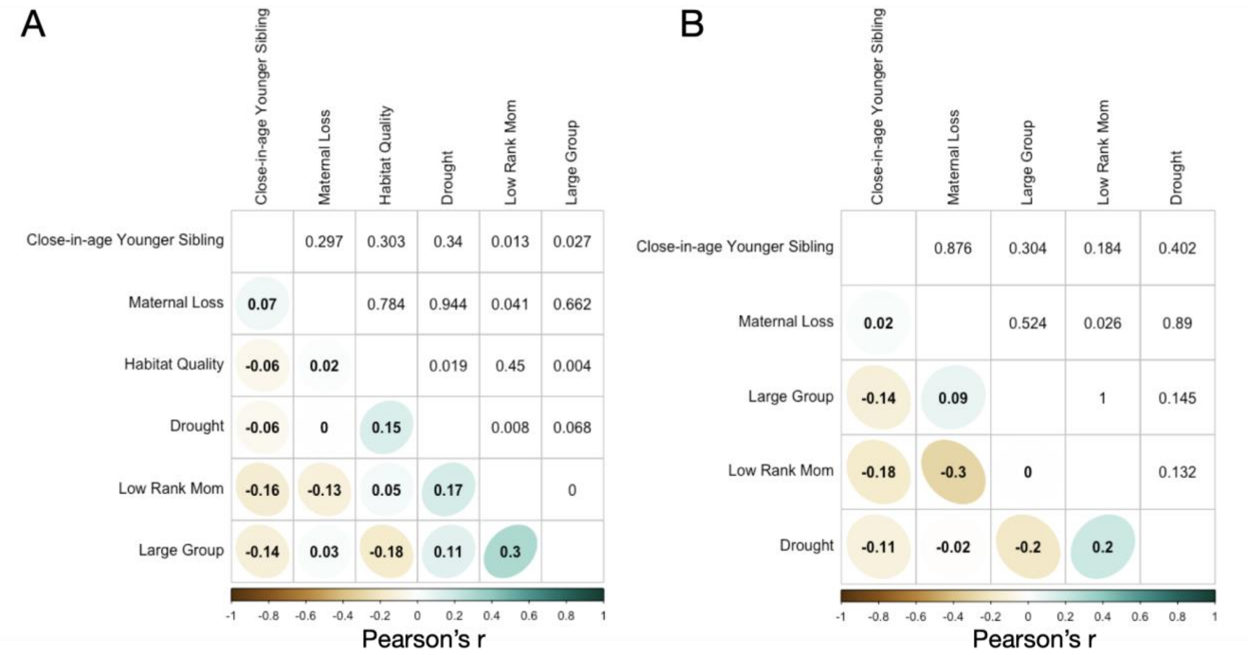


**Figure S2: Pairwise correlations of early life variables across individuals.** (A) Pearson's correlations between exposures to different sources of early life adversity in both the full dataset, and (B) in the subset of individuals born into a low-quality habitat (B). Lower triangle indicates the Pearson's r, colored by the strength of correlation. Numbers in the upper triangle show the p-value for each pairwise correlation.

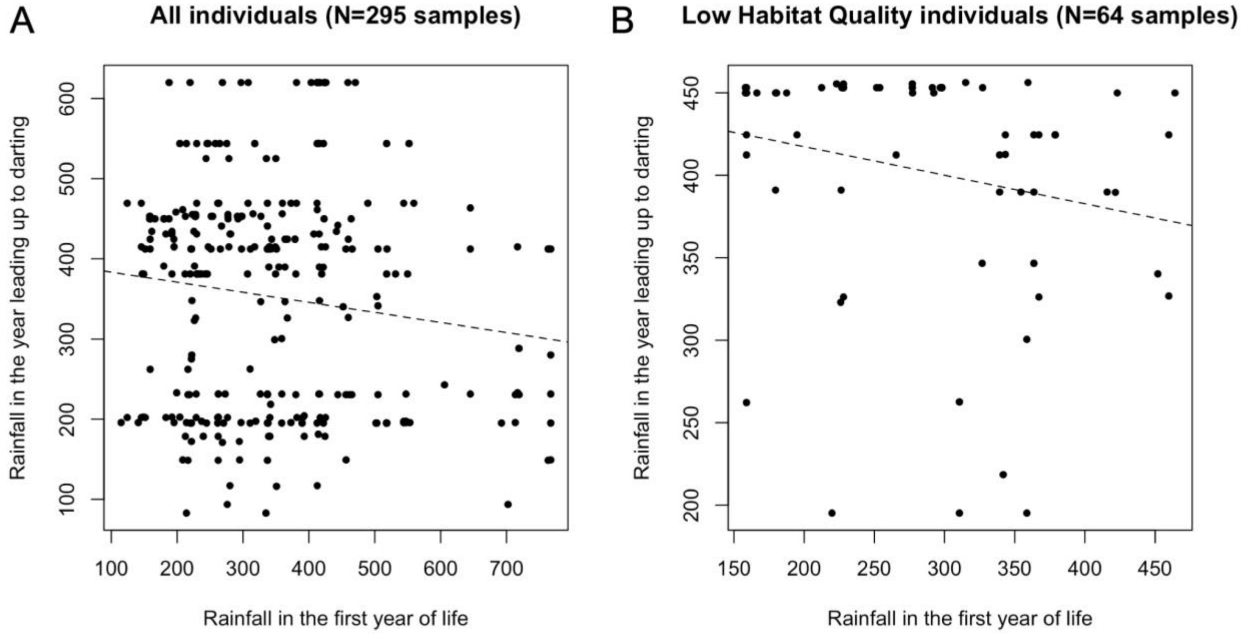


**Figure S3: Rainfall in the first year of life and in the year leading up to darting are weakly negatively correlated.** (A) Cumulative rainfall (mm) in the first year of life (x-axis) versus cumulative rainfall in the year leading up to sample collection in the full dataset ($p=0.02$, Pearson's $R=-0.13$) and (B) in the subset of individuals born pre-habitat shift ($p=0.09$, Pearson's $R=-0.21$).

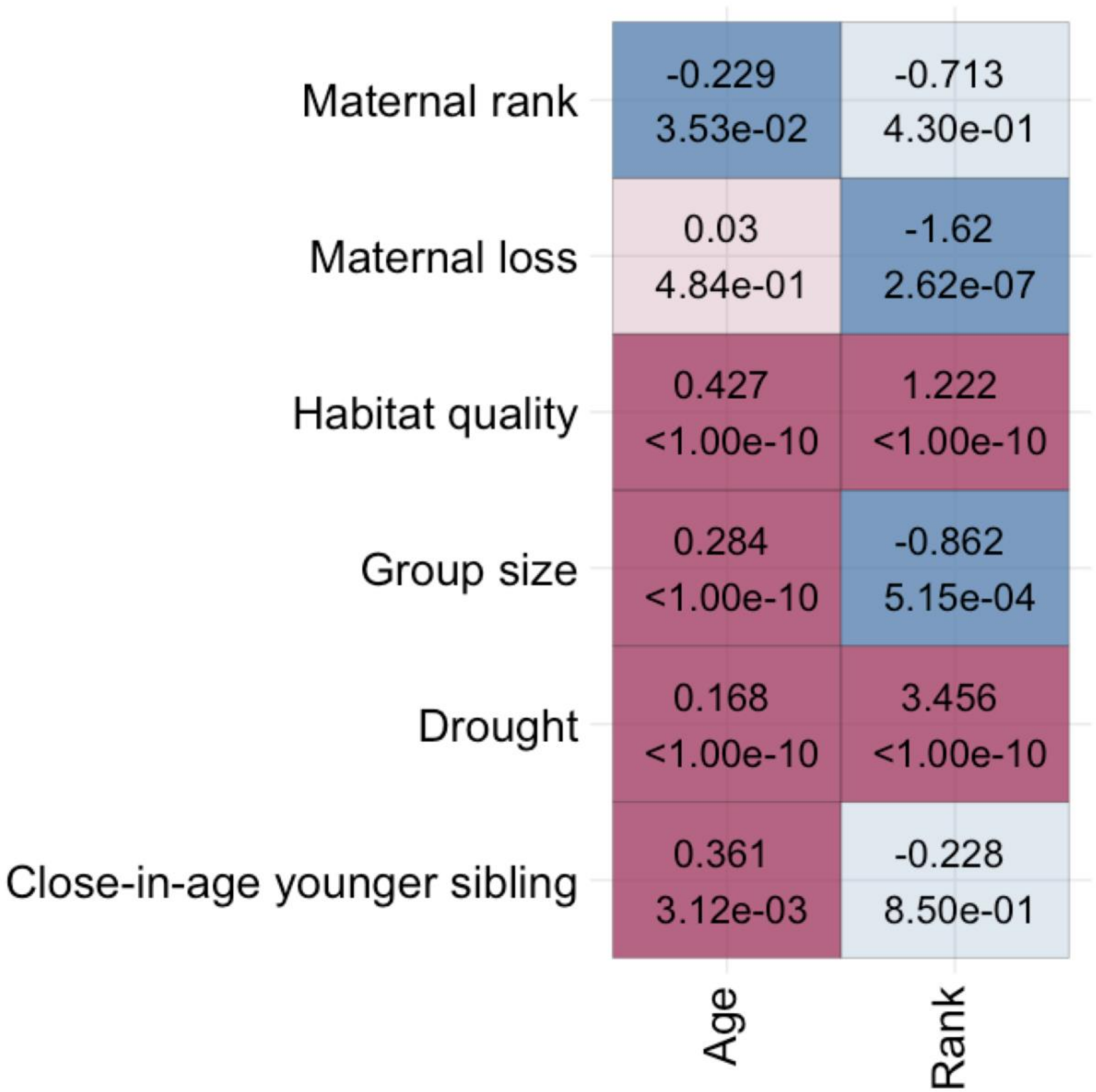


**Figure S4: Overlap between age and rank effects and the effects of early life environment.** Results from Fisher's exact tests for overlap of significant effects of rank or age and each early life variable (10% FDR threshold). Top values show the $\log_2$(odds ratio), lower values indicate p-values. P-values less than 1 x $10^{-10}$ are abbreviated as <1 x $10^{-10}$. Colors indicate the sign of the effect (blue indicates under-enrichment and red indicates enrichment).

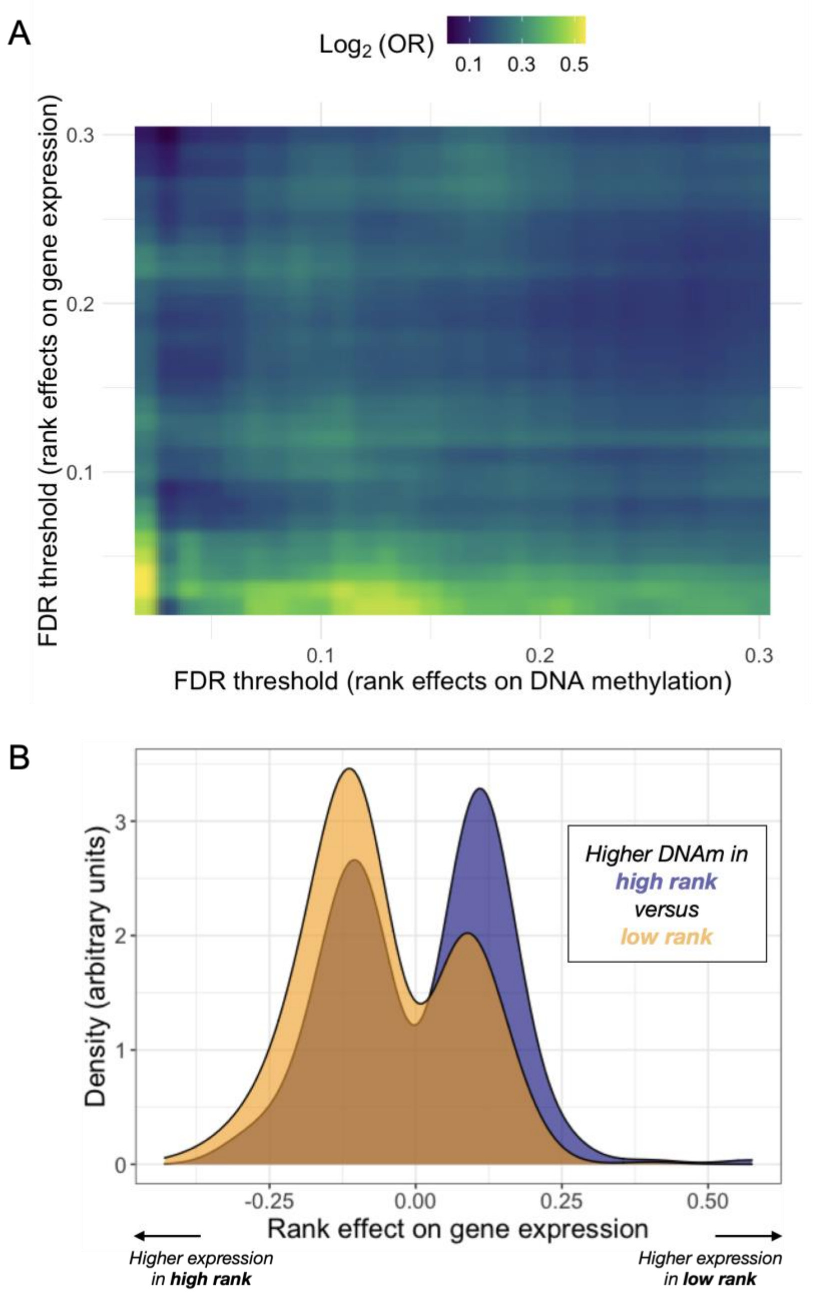


**Figure S5: Dominance rank effects on DNA methylation predict those observed in a separate gene expression dataset.** (A) Overlap between dominance rank effects on DNA methylation within gene bodies (x-axis) and dominance rank effects on gene expression for the same genes (y-axis), across FDR thresholds for discovery. The evidence for overlap increases (yellow colors) with increasingly stringent significance thresholds. (B) Genes containing a rank-associated CpG site exhibit higher methylation levels at that site in low ranking males (orange) when they are expressed more highly in high-ranking males. Higher DNA methylation is observed in high-ranking males (blue) males when those genes are more highly expressed in low-ranking males.
